# *In vitro* Functional Analysis of pgRNA Sites Regulating Assembly of Hepatitis B Virus

**DOI:** 10.1101/2021.09.08.459240

**Authors:** Nikesh Patel, Sam Clark, Eva U. Weiß, Carlos P. Mata, Jen Bohon, Erik R. Farquhar, Daniel P. Maskell, Neil A. Ranson, Reidun Twarock, Peter G. Stockley

**Author notes:** Joint corresponding authors. ^♠^Los Alamos National Laboratory, Los Alamos, NM 87545, USA; & ^°^Electron and Confocal Microscopy Unit (UCCTs), National Centre for Microbiology (ISCIII). Majadahonda, Madrid, Spain.

## Abstract

The roles of RNA sequence/structure motifs, Packaging Signals (PSs), for regulating assembly of an HBV genome transcript have been investigated in an efficient *in vitro* assay containing only core protein (Cp) and RNA. Variants of three conserved PSs, within the genome of a strain not used previously, preventing correct presentation of a Cp-recognition loop motif are differentially deleterious for assembly of nucleocapsid-like particles (NCPs). Cryo-electron microscopy reconstruction of the *T*=4 NCPs formed with the wild-type gRNA transcript, reveal that the interior of the Cp shell is in contact with lower resolution density, potentially encompassing the arginine-rich protein domains and gRNA. Symmetry-relaxation of this reconstruction reveals that such contacts are made at every symmetry axis. We infer from their regulation of assembly that some of these contacts would involve gRNA PSs, and confirmed this by X-ray RNA footprinting. Mutation of the ε stem-loop in the gRNA, where polymerase binds *in vivo*, produces a poor RNA assembly substrate with Cp alone, largely due to alterations in its conformation. The results show that RNA PSs regulate assembly of HBV genomic transcripts *in vitro*, and therefore may play similar roles *in vivo,* in concert with other molecular factors.

## Introduction

Hepatitis B Virus (HBV) has infected over 2 billion people worldwide^1^, ∼240 million of whom are chronically infected after failing to clear an acute primary infection. Within this cohort, failure to suppress the virus leads eventually to liver failure, cirrhosis and cancer, resulting in ∼ 700,000 deaths annually^2^. Despite an effective vaccine, over a million new infections occur every year^3, 4^. For chronically infected patients therapy options are limited. Clinical therapy commonly uses nucleos(t)ide analogue inhibitors of viral polymerase (Pol) but this rarely leads to a cure and elicits rapid resistance mutations^5, 6^. HBV represents one of the largest health challenges of any viral pathogen. A WHO Global Challenge has been established with the goal of making chronic infection treatable by 2030^7^. Novel curative therapies require improved mechanistic understanding of the HBV lifecycle.

The basis of the chronic infection is a covalently-closed, circular DNA (cccDNA) copy of the viral genome which persists in the nucleus as a chromatinized episome^8, 9^. HBV is a para-retrovirus, i.e. a DNA virus that initially packages a positive-sense, single-stranded (ss) pre-genome, pgRNA^10, 11^, into a nucleocapsid (NC) composed of multiple Cp dimers (Fig 1). These form *T*=3 or *T*=4 surface lattices, the latter being dominant. The 3200 bp long genome encodes four overlapping reading frames for polymerase (Pol); surface proteins (HBsAg); the cell regulatory factor protein X; and the core and pre-core proteins (HBcAg and HBeAg, respectively). Pol and Cp proteins are translated from the pgRNA, which also serves as the template for reverse transcription within the NC shell. The pgRNA is a 5’-capped, terminally redundant, poly-A tailed, mRNA transcript ∼3500 nts long.

**Figure 1:**
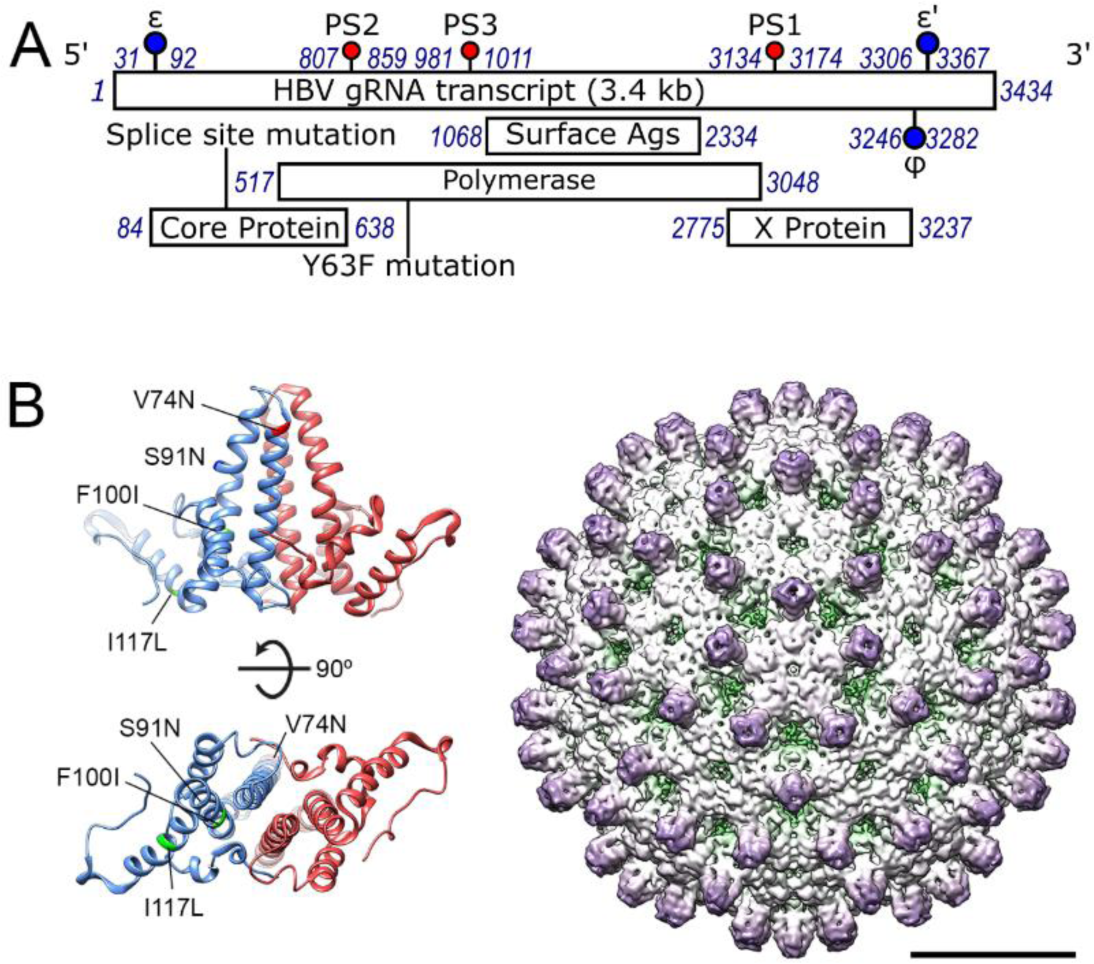
Locations of HBV RNA packaging signals & structure of the *T*=4 NCP. (A) Genetic map of the JQ707375.1 HBV pgRNA with the regions encoding open reading frames shown as bars below. The locations of its PSs, corresponding to homologous sites from NC_003977.1, are shown as lollipops, as are the ε and ϕ sites. Here and throughout, blue numbers refer to cognate genome nucleotide locations. (B) *Left,* Ribbon diagrams showing amino acids 1 to 143, of the Cp dimer, with the monomers coloured red and blue (PDB ID 3J2V) viewed from the side with the interior of the particle below (top) or from below (bottom). The locations of the amino acid changes between strains NC_003977.1 and JQ707375.1 are indicated on the blue monomers. *Right,* I1 reconstruction at ∼4.7 Å resolution of the NC_003977.1 *T=*4 HBV NCP (EMD-3715) reassembled around an oligonucleotide encompassing its PS1, viewed along a five-fold axis (bar = 100 Å).^12^ The projections correspond to the helices in the Cp dimer, left.

Previously, RNA SELEX^12, 13^ against recombinant full-length HBV Cp (183 aa long) dimers from strain NC_003977.1 was used to isolate aptamers whose sequences align with genomic sequences in the cognate pgRNA. Analysis of these genomic matching sites identifies their common features. Each site has the potential to form a stem-loop with a defined loop sequence motif, -RGAG-. Such sites are highly conserved across the pgRNAs of many strain variants. These HBV sites, as isolated RNA oligonucleotides, trigger *in vitro* sequence-specific NCP formation at nanomolar concentrations^12^. We propose that they act as Packaging Signals (PSs) helping to ensure faithful encapsidation of the pgRNA, by analogy with the mechanism regulating assembly of many *bona fide* ssRNA viruses^14–20^.

Here we show that homologous PSs occur in similar genome locations in a commercially-available strain variant (JQ707375.1), which was not included in the previous analysis. These sites regulate *in vitro* assembly of *T*=3 and *T*=4 nucleocapsid-like particles (NCPs) in the context of a long genomic gRNA fragment lacking a 5’ cap and a poly-A tail. Regulation occurs at least in part through Cp recognition of the conserved loop –RGAG-motif of the PSs. Some of these contacts remain in the assembled particle. Symmetry expansion^21–24^ of an icosahedrally-averaged ∼3.2 Å resolution cryo-electron microscopy (cryo-EM) reconstruction of the reassembled *T*=4 NCPs reveals multiple contacts at all symmetry axes between encapsidated density (the C-terminal arginine-rich domains (ARDs) of Cp’s & gRNA) and the globular Cp shell. X-ray RNA footprinting (XRF)^25–27^ confirms that the most conserved PS makes one such contact via its –RGAG-motif. Assembly *in vitro* occurs in the absence of viral polymerase which binds a stem-loop (ε) on the pgRNA^28^, and post-translational modifications of the Cp. Both features assist NCP formation *in vivo*^28–32^. Mutation to prevent ε interacting with the distal ϕ site creates a genomic fragment that is considerably larger than both wild-type and PS mutant RNAs, as well as being a poor *in vitro* assembly substrate. These data highlight that PS-mediated assembly via the formation of multiple PS-Cp contacts promotes NCP-like formation, in the absence of other mechanisms regulating assembly.

## Results

### Identification of PS sites in HBV strain JQ707375.1

Evolutionarily conserved PS sites in the pgRNA of NC_003977.1 (subtype *ayw*) were identified by aligning anti-Cp RNA aptamers against the sequences of 16 strain variant HBV pgRNAs, chosen at random from the ∼750 sequences then available in GenBank. Mfold^33^ suggests that each of the matched sites is potentially able to fold into a stem-loop with an over-represented sequence, 5′-RGAG-3′, in the loop^12^. The three most highly conserved NC_003977.1 sites, PSs1-3, as oligonucleotides, trigger sequence/structure-specific *in vitro* assembly of Cp, mostly into *T*=4 NCPs. Sequence alignment and Mfold^33^ readily identifies putative PS homologues of these sites in the JQ707375.1 RNA sequence (Fig 2; Materials & Methods). Each of the JQ707375.1^34^ PS homologues is predicted to fold into a stem-loop, the latter presenting a purine-rich tetra-nucleotide motif. PSs 1 & 3, have ideal consensus Cp-recognition motifs, -GGAG-& -AGAG-, respectively, whilst the homologous PS2, has a slight variation (-GAAG-). Their secondary structures and stabilities vary, as expected^12^.

**Figure 2:**
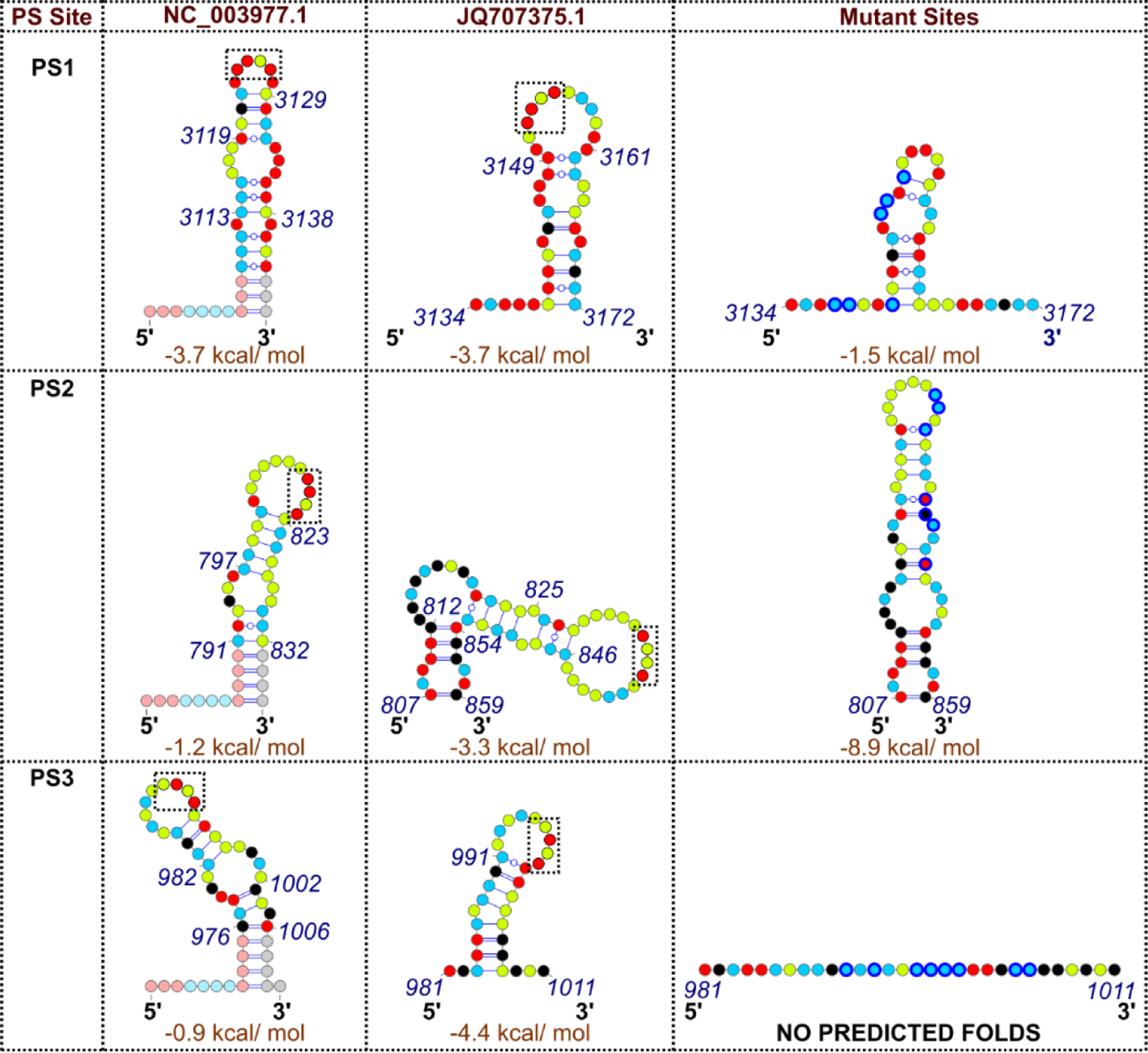
Homologous PSs in the two HBV strains and their variants. HBV NC_003977.1 and JQ707375.1 PSs1-3, and their variant, sequences and Mfold secondary structures^12^. Faded nucleotides represent G:C clamps and 5’-leaders used to force the NC_003977.1 sequences to adopt single folds and to facilitate dye-labelling^12^. Here and throughout, RNA nucleotides are shown as coloured dots (*green - A, black - C, red - G, blue - U*), Watson-Crick base pairs are indicated as lines, which are interrupted by circles for G-U pairs. The predicted folding free energies of the unliganded RNA fragments are also shown. These do not include the non-HBV sequences. -RGAG-motifs are shown boxed (black), and mutated nucleotides within PS variants are outlined in dark blue.

To investigate the PS-mediated NCP assembly hypothesis, and determine the roles of PSs in assembly in the context of the genome, we used a transcript assembled from the commercially available strain JQ707375.1^34^. This was isolated from a patient with a lamivudine-resistant infection and is therefore thought to be replication competent. The ∼3400 nt assembled RNA is closer to the size of full-length pgRNA but lacks the 5’ cap and 3’ polyadenylation. It shares 90.7% nucleotide sequence identity with the same region of NC_003977.1, from which the recombinant Cp used in assembly assays was expressed^12^. There are just 4 amino acid sequence changes between the Cp’s of these strains: (NC_003977.1 vs. JQ707375.1, respectively) V74N; S91N; F100I & I117L, all of which lie within the first 149 amino acids of core (Fig 1B) which form the outer portion of the NCP shell. None would be expected to interact with the pgRNA. NC_003977.1 Cp was therefore used in the *in vitro* assembly assays with the JQ707375.1 transcript.

### In vitro assembly with the JQ707375.1 RNA transcript

PS-mediated assembly of ssRNA viruses at nanomolar concentrations^12, 16, 35^ mimics the genome packaging specificity outcomes of natural infections^36^. These conditions differ from most *in vitro* reassembly studies which are usually carried out at much higher concentrations^37, 38^. Previously, PS oligonucleotide-induced HBV NCP assembly was monitored using single-molecule (sm)fluorescence correlation spectroscopy (smFCS) with dye-labelled RNA oligonucleotides^12, 16, 39^. These assays monitor real-time changes to the hydrodynamic radius (R*_h_*) of the labelled RNA, and end-products were analysed by negative stain EM (nsEM). However, the small sample sizes of such experiments prevent more detailed analysis. To overcome that limitation here, NCPs were reassembled in 96-well plates under sm conditions using a liquid-handling robot (Fig 3A). These samples were pooled and concentrated (∼10 fold) to allow subsequent fractionation and further analysis. Well over 90% of the input RNA was recovered from this step, presumably with the rest lost in the concentrator. Subsequent analysis assumed that the material remaining soluble reflects the endpoint of each titration.

**Figure 3:**
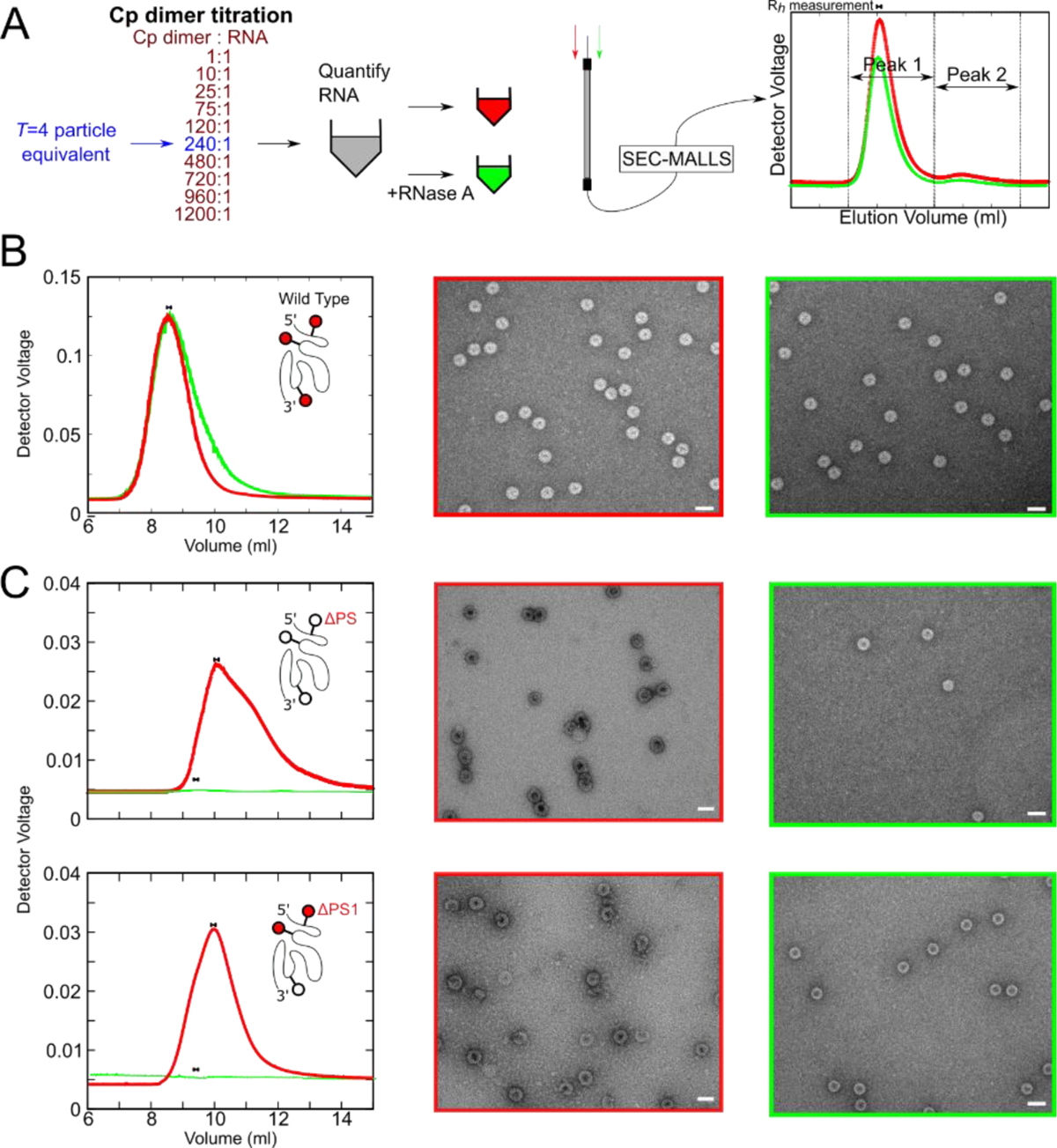
Roles of PSs in regulation of *in vitro* assembly of HBV NCPs. (A) Cp dimer was titrated at 10 min intervals (left) into heat-annealed (Methods) HBV gRNA transcript in a 96-well plate. Reactions were then pooled (grey) and split into 2 (red/green), one of which was treated with RNase (green). Aliquots of each were visualised by nsEM and the remaining sample analysed by SEC-MALLS chromatography, recording the elution volumes (graph, right) and hydrodynamic radii (R*_h_*, bar above peak) of particulate reassembly products from their light-scattering (Methods). (B) *Left*: LS traces of the reassembly products with PS variants of 1 nM HBV gRNA and HBV Cp dimer: red and green traces are pre- and post-RNase treatment, respectively. *Middle / Right*: nsEMs of the assembly products shown (left). Scale bars = 50 nm. (C) *Left:* LS traces, as in (B), with 1 nM ΔPS (top) and ΔPS1 (bottom), and *Middle / Right*: nsEMs of the assembly products shown.

Titrations of Cp dimer, prepared from recombinant NCPs using 1.5 M guanidine hydrochloride (GuHCl), comprising 10x 2 μL aliquots from one of six different stock concentrations, were made into an initial (180 μL) solution containing gRNA (1.1 nM). These dilute the gRNA to a final concentration of 1 nM, whilst spanning sub-stoichiometric to a 4-fold molar excess for formation of a *T*=4 NCP around each gRNA. Each titration point was equilibrated for 10 min at room temperature (∼20 °C) before addition of the next aliquot. This titration allows assembly initiation to occur at the highest affinity PSs on each gRNA molecule and then proceed to completion^37^, and yields are >80% as judged by input gRNA. The final molar excess of Cp dimer ensures that any assembly defects detected are not a consequence of non-functional Cp dimers. Assembled NCPs appear as separated, negative-stain excluding particles in TEM, with A_260/280_ ratios consistent with each particle containing a single, full-length copy of the transcript (see below).

The products from the titration reactions were analysed by absorption spectroscopy and the relative efficiency of complete NCP formation probed by treatment of one of half of the sample with RNase A. Both aliquots were then fractionated by size-exclusion chromatography with the eluting material detected via multi-angle, laser light-scattering (SEC-MALLS). Light-scattering peaks were collected, re-concentrated to ∼1 mL, and their absorbance values re-measured. The SEC-MALLS signals were used to estimate assembly efficiency and the hydrodynamic radii (R*_h_*) of eluting species (Figure 3A, Table 1).

**Table 1:** Characterisation of NCPs reassembled around gRNA constructs. All values are post-nuclease treatment unless stated.

| **gRNA** | **A<sub>260</sub> ( Before and after RNase A addition)** | **A<sub>260/280</sub>** | **RNA yield (%)** | **R<sub>h</sub> / nm** |
| --- | --- | --- | --- | --- |
| Wild-type | 0.12 → 0.097 | 1.49 | 81 | 18.2 |
| ΔPS | 0.019 → 0.0067 | 0.91 | 35 | 6.2 |
| ΔPS1 | 0.048 → 0.004 | 1.07 | 8 | 9.2 |
| ΔPS2 | 0.128 → 0.095 | 1.55 | 74 | 18.4 |
| ΔPS3 | 0.119 → 0.069 | 1.3 | 58 | 17.2 |

For JQ707375.1 RNA transcripts the reassembly products prior to RNase treatment (Fig 3B) elute as a single, symmetrical peak, ∼8.5 mL after application to the column. Unassembled Cp dimer, present from the titration, is much smaller than NCPs and invisible on these plots. RNase-treated aliquots elute essentially identically with a very similar yield, although their peak has a slight low-side shoulder. Prior to chromatography, both aliquots contain identical stain-excluding particles in nsEM, consistent with *T*=4 NCPs. Their apparent R*_h_* values (∼18-20 nm) match those for NCPs formed by Cp recombinant expression in *E. coli* (∼19 nm, Sup Fig 1), and are similar to values of ∼25 – 32 nm determined by previous cryo-EM and smFCS measurements, respectively. The results show that NC_003977.1 Cp dimer successfully reassembles nuclease-resistant *T*=4 NCPs around JQ707375.1 gRNA (Table 1), validating the assumption about the common functionality of Cp’s between HBV strains.

### Probing the role(s) of PSs1-3 in in vitro assembly of gRNA

Sequence variants were designed to investigate the role(s) of PSs in regulating assembly with JQ707375.1 transcripts in the *in vitro* assembly assay. Variants remove loop recognition motifs and/or destabilize the secondary structures of the stems presenting the motif, whilst retaining wherever possible the global gRNA fold, as predicted by Mfold^33^. PS1 lies within an especially G-rich (50%) region and consequently encompasses several potential alternative -RGAG-motifs. This made subtle motif ablation impossible. Instead, six G to U mutations were introduced into the 5′ leg of the PS stem destabilising it (predicted free energy of folding going from −3.7 to −1.3 kcal mol^-1^), and increasing the bulge size, whilst moving the –RGAG-motif to a more central position within the loop (Fig 2). For PS2 and PS3, purine to U mutations can be introduced directly into the loops, replacing their recognition motifs -GAAG- and -AGAG-, with -UUAU- and -UUUU-, respectively. It was also possible to destabilize the base-pairing in their stems via the following substitutions: in the 3′ leg of the PS2 stem-loop (-UUAAAAUUA- to -UAGCUUUG- (nts 841-848)), and either side of the loop in PS3 (-AUAUAUUUUGGGAA- to -UUUUAUUUUGGCUU- (nts 992-1005)). These substitutions prevent formation of alternate -RGAG-motifs in either PS. Disrupted PS2 is predicted to fold as a very stable (−8.9 kcal mol^-1^) stem-loop, with a multiply interrupted stem. The mutations within PS3 ablate all possible secondary structures using the default Mfold parameters (Fig 2). The global folding effects of these changes were estimated by folding a 300 nt region centred on each PS using a sliding 60 nt window noting the frequency of defined structures. This procedure identified no long-range base-pairing issues and confirmed the greatly reduced occurrence of a Cp-recognition motif in a loop in the PS1 variant, together with the complete ablation of PS2 and PS3 recognition signals and secondary structures (data not shown).

The effects of the variant PSs on NCP assembly were assessed in transcripts containing all three variant PSs (ΔPS), or with individual variants (ΔPS1, ΔPS2 and ΔPS3) (Fig 3C, Table 1). ΔPS gRNA does not assemble *T*=4 NCPs significantly under these conditions. Its pre-RNase assembly product elutes later than a *bona fide* NCP (10 mLs after being applied to the column) and has a pronounced low-side tail. In nsEM, it appears to produce small quantities of malformed and aggregated particles which are readily penetrated by stain. Absorbance measurements are consistent with this, the peak containing only ∼10% of the wild-type amount of RNA. Particle formation, however, remains RNA-dependent since RNase treatment eliminates all light-scattering material, revealing only a small number of apparently correctly assembled particles. This result confirms the assumption that these sites act as PSs within JQ707375.1 gRNA. Alteration of only 24/3200 nucleotides in this RNA transcript completely prevents assembly. Since it is the same length as the wild-type sequence, assembly cannot rely purely on favourable electrostatic interactions, as has been proposed for many single-stranded (ss) RNA viruses^40–45^, i.e. it implies assembly regulation by the RNA. The sequence variations in ΔPS gRNA do not alter its hydrodynamic radius significantly (Sup. Fig 2) confirming that this regulation is based on sequence-specific recognition of PS sites.

ΔPS1 pgRNA also assembles poorly compared to the wild-type (Fig 3C, Table 1), eluting (∼9.5 mL vs ∼8.5 mL for WT) as a roughly symmetrical peak. The material before RNase treatment consists mostly of separate but misshapen particles of widely differing radii, which are mostly freely penetrated by negative stain. As with the ΔPS variant, assembly is dependent on the RNA, with RNase treatment eliminating all light-scattering material, although there are many more particles roughly the size of an NCP than with the ΔPS variant transcript. They are all freely penetrated by negative stain. These results arise from variation of just 6 nucleotides across the gRNA, and an –RGAG-motif is still present in the smaller loop of the variant. Since the ΔPS and ΔPS1 transcripts have distinct assembly properties, PSs 2 & 3 must also contribute differentially to assembly regulation. Analysis of their individual reassembly reactions suggests, however, that these are less significant than for PS1 (Sup Fig 3). Both variants produce products that co-elute with *bona fide* NCPs assembled around wild-type gRNA. They elute as symmetrical peaks containing mostly stain-excluding, single particles. Their hydrodynamic radii are also very close to that of wild-type (∼17-18 nm vs ∼18 nm), although their susceptibility to RNase (Table 1) implies that they fail to form completely closed shells. ΔPS3 appears more deleterious in terms of closed shell formation than ΔPS2.

These results are consistent with a PS-mediated *in vitro* assembly mechanism for the HBV JQ707375.1 gRNA. Multiple PSs within the genomes of ssRNA viruses^15, 46, 47^ vary around a consensus sequence defining a hierarchy of binding affinities for cognate capsid proteins. This preferred assembly pathway around cognate viral RNA prevents multiple assembly initiation events occurring on the same RNA, ensuring efficient capsid assembly, and avoiding the formation of off-pathway kinetic traps^37^. It implies that PSs act cooperatively but differentially during assembly, as seen here. This mechanism creates a fixed spatial relationship between the pgRNA and Cp shell, and for several RNA bacteriophages these interactions have been revealed directly by asymmetric cryo-EM reconstruction^48, 49^. Zlotnick and colleagues ^50^ have proposed that a preferred pgRNA conformation within the NCP would facilitate its reverse transcription. To investigate that possibility, we determined the structure of the NCP assembled around the wild-type RNA transcript.

### The gRNA containing NCP has ordered internal density

A cryo-EM reconstruction was calculated using particles of purified NCPs containing the HBV JQ707375.1 RNA transcript (Methods & Table 2). After data collection, 127,410 *T*=4 (84%) and 23,257 *T=*3 (16%) particles were chosen for further analysis. The ratio of *T*=4:*T*=3 particles in this dataset is similar to that obtained following reassembly around the NC_003977.1 PS1 oligonucleotide^12^. 2D classification reveals that most *T*=3 particles and ∼18% of the *T*=4 particles were heterogeneous, with many particles appearing empty. No further image processing was performed on these. A homogenous dataset of the remaining *T*=4 particles was subjected to icosahedrally-averaged refinement yielding a ∼3.2 Å resolution map. This reconstruction reveals a Cp layer similar to those seen in previous EM and crystal structures. Density for Cp chains is complete up to the start of the ARD (Fig 4, Sup Fig 4)^51–54^. The four Cp monomers of the asymmetric unit were built into the map using a previous structure of the NC_003977.1 Cp^54^ (PDB 3J2V) as a starting model. At this resolution, the four distinct Cp monomers of the quasi-equivalent capsomer, chains A-D, can be built using polypeptide chains 144, 143, 144 and 147 residues long, respectively (Sup Figs 4 & 5). The icosahedrally-ordered Cp surface lattice surrounds a second, less-ordered internal layer of density, which may well include density for the encapsidated gRNA transcript and the Cp ARD domains.

**Figure 4:**
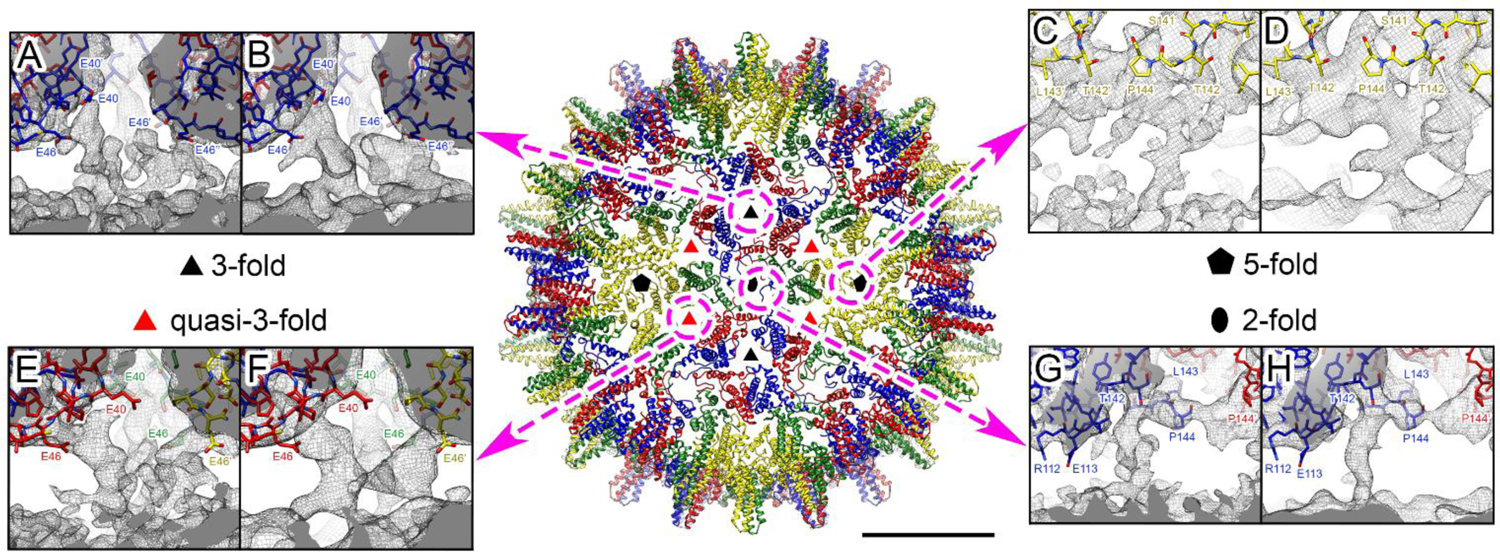
Cryo-EM reconstruction of the *T*=4 NCP formed with JQ707375.1 gRNA. *Centre:* Atomic model of the front-half of the HBV *T*=4 NCP shown as ribbon diagrams, coloured in yellow (A monomer), green (B monomer), blue (C monomer) and red (D monomer) for the quasi-equivalent monomers, and viewed along a two-fold axis built into the icosahedrally-averaged cryo-EM density map at 3.2 Å resolution (bar = 100 Å). Symbols indicate icosahedral symmetry axes. (A-H) Dashed (pink) circles indicate the locations of individual NCP symmetry axes, including the quasi-3-fold, and the arrows lead to boxes showing the transiting internal density associated with each of these locations. Cryo-EM density maps are shown as a grey mesh with the centre of the NCP towards the bottom. Cp atomic models are shown as sticks. Amino acid side-chains adjacent to the internal layer of density are indicated and coloured by heteroatom. The atomic model is fitted into the non-filtered shell of the Class 3 density map obtained after symmetry expansion and focused classification (A, C, E, G), and for clarity into the same map low-pass filtered to 5 Å (B, D, F, H), both shown at 1.5 σ. (A, B) Contacts located at three-fold vertices involve the E40-C48 sequence in Cp (including the E40-C43 alpha-helix and the S44-C48 loop) of three C monomers. The side chains of residues E40 and E46 are directly pointing to the internal density shell. (C, D) Contacts located at five-fold vertices involve five A monomers (pentamer). (E, F) Contacts located at quasi-three-fold vertices involve the E40-C48 sequence in Cp (including the E40-C43 alpha-helix and the S44-C48 loop) of one A monomer, one B monomer and one D monomer. The side chains of residues E40 and E46 point directly at the internal desnity shell. (G, H) Contacts located at two-fold vertices involve two C monomers and two D monomers (hexamer).

**Figure 5:**
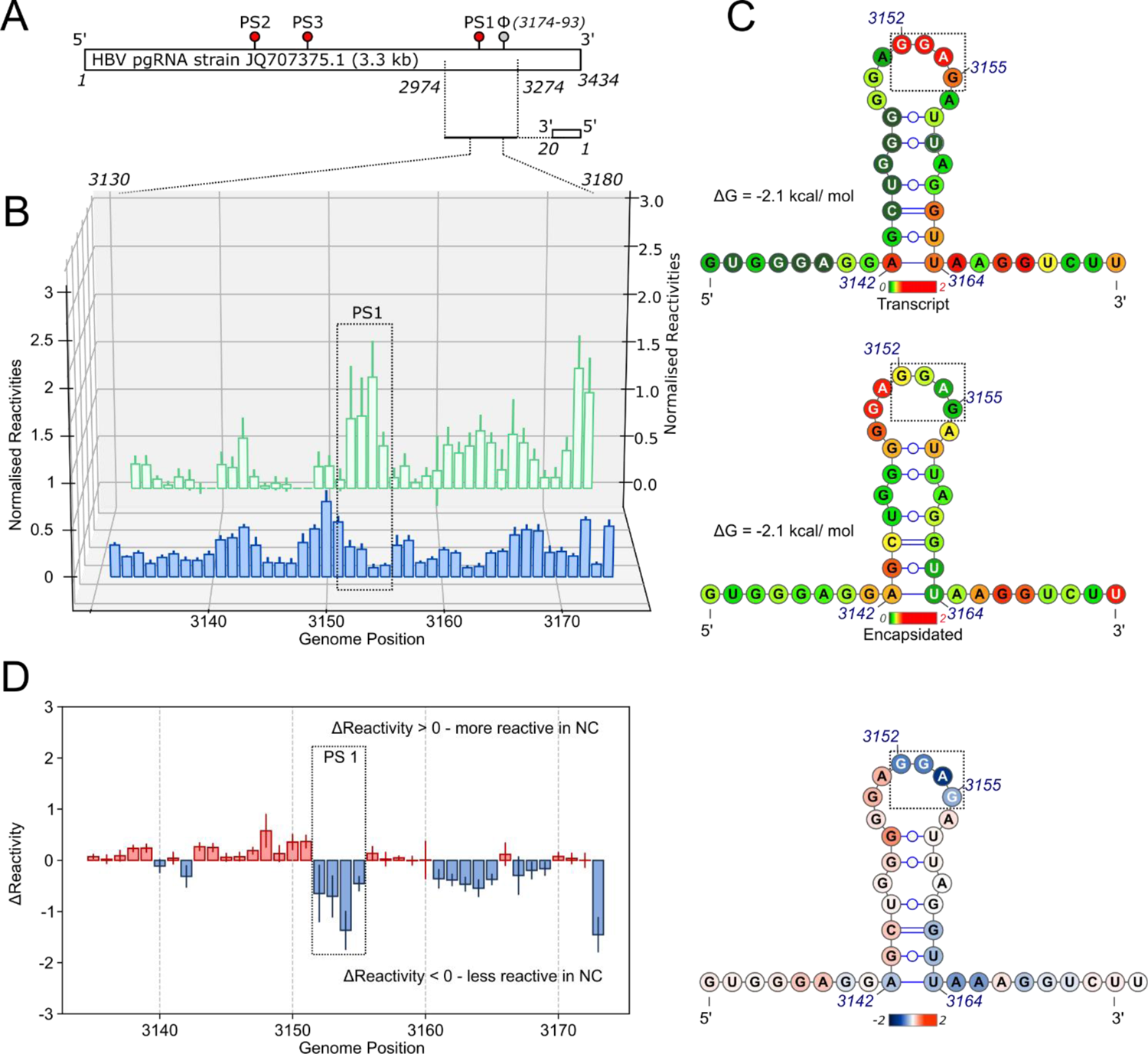
RNA footprints of the PS1 region. (A) Protein-free transcripts of the gRNA, or following reassembly and purification of NCPs, were hydroxyl radical footprinted as frozen samples by exposure to X-rays.^25, 26, 58^ RNAs were recovered from frozen samples. Reverse transcription from a 5’fluorescently-end-labelled primer complementary to the pgRNA 3’-end was used to assess nucleotide reactivities in a fragment encompassing PS1 using capillary electrophoresis. (B) Waterfall plot of the per nucleotide reactivities and associated standard errors from triplicate samples of transcript (green) or NCP (blue). (C) Secondary structures of the PS1 folds showing their XRF reactivities. Nucleotides are coloured red to green depending on their relative reactivity on exposure to the beamline (red → green = reactive → inert (reactivity of 2.0 → 0)), scale shown below each fold. (D) *Left:* The average reactivity at each nucleotide across the PS1 sequence is shown. Red/blue indicate positions more/less reactive in the NCP. *Right:* Average reactivity at each nucleotide shown upon the secondary structure of PS1. Nucleotides are coloured blue/red depending on their reactivity within the NCP (blue → red = less → more reactive (Δreactivity of −2.0 → 2.0)), scale shown below the PS1 fold.

**Table 2:** Cryo-EM structure. Cryo-EM data collection, refinement and validation statistics.

|  | **HBV NCP sym I1**<br>**EMD-11700 / PDB:**<br>**7ABL** | **HBV NCP sym exp**<br>**EMD-11702** | **HBV NCP sym C1**<br>**EMD-11701** |
| --- | --- | --- | --- |
| **Data collection and processing** |  |  |  |
| Microscope | FEI Titan Krios | FEI Titan Krios | FEI Titan Krios |
| Detector | FalconIII (linear mode) | FalconIII (linear mode) | FalconIII (linear mode) |
| Magnification | 75,000x | 75,000x | 75,000x |
| Voltage (kV) | 300 | 300 | 300 |
| Electron exposure (e<sup>-</sup>/Å<sup>2</sup>) | 53.1 | 53.1 | 53.1 |
| Exposure per frame (e<sup>-</sup>/Å<sup>2</sup>) | 0.9 | 0.9 | 0.9 |
| Defocus range (μm) | -0.75 to -2.5 | -0.75 to -2.5 | -0.75 to -2.5 |
| Pixel size (Å) | 1.065 | 1.065 | 1.065 |
| Micrographs collected (no.) | 2,658 | 2,658 | 2,658 |
| Initial particles (no.) | 150,667 | 104,337*60 =<br>6,260,220 | 150,667 |
| Final particles (no.) | 104,337 | 1,254,013 | 104,337 |
| Symmetry imposed | I1 | C1 | C1 |
| Map resolution (Å) | 3.2 | 3.5 | 4.2 |
| FSC threshold | 0.143 | 0.143 | 0.143 |
| Map resolution range (Å) | 3.02 – 3.59 | 3.3 – 6.7 | 3.7 – 7.2 |
| **Refinement** |  |  |  |
| Model resolution (Å) | 3.21 |  |  |
| FSC threshold | 0.5 |  |  |
| Mask correlation coefficient | 0.87 |  |  |
| Map sharpening B factor (Å<sup>2</sup>) | -158 |  |  |
| Model composition |  |  |  |
| Non-hydrogen atoms | 277,380 |  |  |
| Protein residues | 34,680 |  |  |
| ADP (B-factors)<br>min/max/mean |  |  |  |
| Protein | 9.60/85.84/27.30 |  |  |
| R.m.s. deviations |  |  |  |
| Bond lengths (Å) | 0.007 |  |  |
| Bond angles (°) | 0.857 |  |  |
| Validation |  |  |  |
| MolProbity score | 1.30 |  |  |
| Clashscore | 2.57 |  |  |
| Rotamer outliers (%) | 0.58 |  |  |
| Ramachandran plot |  |  |  |
| Favored (%) | 96.14 |  |  |
| Allowed (%) | 3.68 |  |  |
| Outliers (%) | 0.18 |  |  |

Reconstruction of the same dataset without the imposition of symmetry fails to reveal any further molecular detail at 4.2 Å resolution, the internal density appearing as a mostly continuous layer underneath the Cp shell (Sup Fig 6). To reduce the impacts of the Cp layer on the reconstruction, the signal from the globular region of the Cp was subtracted from the particles in the same dataset, and a 3D classification by alignment of the remaining density calculated revealing five similar classes (not shown). The icosahedrally-averaged reconstruction of the *T*=4 particle was then symmetry expanded^21–24^ following removal of much of the Cp shell using a spherical mask. The remaining density was subjected to further 3D classification without alignment into a further five roughly equally populated classes (Sup Fig 7A). These were reconstructed without imposing symmetry^21, 55^, yielding maps with global resolutions of 3.5 – 3.6 Å (Sup Fig 7). The maps were low-pass filtered to 5 Å resolution for clarity revealing an asymmetric cage-like density under the Cp shell at this lower resolution, presumably corresponding to either the C-terminal ARDs or gRNA, or both, in close contact with the Cp shell (Fig 4, Sup Fig 7). Superposition of the internal density obtained by asymmetric reconstruction of the NCP assembled around NC_003977.1 PS1 (EMD-3714), into this map shows that this feature, which we presume includes multiple PS RNA oligonucleotides^12^, overlaps with the internal density seen with the NCP gRNA. Additional refinement failed to resolve these features further here, perhaps due to conformational heterogeneity or flexibility^56, 57^.

Interestingly, these maps reveal multiple fingers of density at every particle vertex that transit the gap between the Cp layer and the internal density. Such bridges, illustrated in Fig 4 based on the Class 3 data (Sup Fig 7D) could be the result of the formation of multiple Cp-gRNA contacts in the NCP, i.e. as expected for PS-mediated assembly. These density bridges 5-fold and 2-fold positions (Figs 4C, D and G, H, respectively) appear to be different at the different symmetry axes. For both, the internal density fuses with that of the Cp subunits adjacent to residue Pro144, i.e. just before the start of the ARD domain, and could represent extensions of the polypeptide chain at these points. The bridging contacts at the 3-fold (Fig 4A, B) and quasi-3-fold vertices (Fig 4E, F) are much more similar to each other but distinct from those at 2- and 5-fold axes. They fuse into the density of the Cp around residues E40 and C48. Both these amino acids are in well-ordered sections of the globular fold and there is no unassigned Cp density in this part of the map.

In order to test the idea that there are RNA PS-Cp contacts in the reassembled NCP, we used XRF^27, 58, 59^. To see if PS1, the most important PS within the JQ707375.1 transcript for regulating assembly *in vitro* (Fig 3C) contacts Cp in the reassembled NCP, XRF nucleotide reactivities across a genomic region encompassing PS1 were determined for both NCP and free transcript (Methods). Both reassembled NCP and transcript were flash-frozen and exposed to the X-ray beam on the customised beam-line at the NSLS-II in Brookhaven for various times ranging from 0-100 milliseconds^60^. Samples were returned to the host laboratory in the frozen state and RNA extracted/prepared for primer extension with a dye-labelled DNA primer annealed at nucleotide positions 3308-3328 (Fig 5). The reverse transcription extension products were then analysed using capillary electrophoresis (CE), and a combination of published and in-house software^27, 61^. Reassembled and protein-free gRNA is predicted to adopt the same stem-loop structure at PS1. This fold places the –RGAG-recognition motif (boxed in Fig 5C) in the 3’ half of the loop, as expected (Fig 2). The reactivities of the nucleotides in these two RNA stem-loops are shown colour coded (dark green to red corresponding to unreactive to highly reactive, respectively), and as a difference map in Fig 5 D. Nucleotides within the loop show the most significant differences in reactivity between the two states. In the transcript, the nucleotides of the -RGAG-motif have high reactivities (red), with the 5’ and 3’ neighbouring nucleotides being much less reactive. In the NCP, however, these reactivity patterns across the motif are largely reversed. All four nucleotides of the recognition motif become largely unreactive, whereas their 5’ and 3’ neighbours become more reactive, an effect that extends into the uppermost base pair of the stem (Fig 5B, C). These data are consistent with a direct interaction between the NCP Cp layer and the PS1 recognition motif. There are also slight conformational rearrangements in the loop in the NCP resulting in increased flexibility of the non-contacted nucleotides.

*In vivo* NCP assembly is known to be affected by Cp phosphorylation and/or the binding of replicase at the ε site^28–32^, both of which are missing in the *in vitro* assay. The JQ707375.1 transcript encompass both ε and ϕ sites (Fig 1), and these sites are thought to form long-range base pairs making the RNA topology important for assembly^62, 63^. In order to test this idea, we carried out reassembly with a variant RNA transcript of its ε site (Sup Fig 2) preventing it from base-pairing with the ϕ site. Strikingly, this variant gRNA, unlike those carrying the PS mutations, has a significantly increased hydrodynamic radius (27± 0.4 *vs*. 17 ± 0.3 nm, respectively). It is also a poor *in vitro* assembly substrate, although not as poor as the ΔPS and ΔPS1 variants, implying that *in vitro* and presumably also *in vivo*, both RNA topology and PS-Cp contacts contribute to assembly efficiency. It has been argued that ε is the most important pgRNA sequence since *in vivo* it is possible to encapsidate heterologous RNAs apparently encompassing only this site from the HBV genome. However, in these experiments the ε site was flanked by additional HBV genomic sequences which sequence/S-fold analysis (Sup Info.) suggests contain multiple unrecognised PS sites akin to those analysed here. In addition, in a fusion of this fragment to a 3’ *lacZ,* an assumed non-specific heterologous RNA construct, an additional PS site is added. The combinatorial effects of such sites would explain their packaging, ε sites being packaged whilst *lacZ* alone is not (Sup Tables 1 & 2).

## Discussion

The assembly of infectious HBV is a complex process that many people have studied *in vivo* in suitable cell culture systems. Here we have used a minimised molecular system to investigate whether the RNA stem-loops identified previously stimulate *in vitro* assembly of an RNA transcript encompassing most of the coding region of a strain not previously part of our analysis (JQ707375.1). The results establish that in such an assay this gRNA transcript is efficiently encapsidated. Modification of the gRNA to prevent or alter presentation of the Cp-recognition sequence (-RGAG-), at three putative PS homologues of sites studied previously as oligonucleotides, results in varying assembly deficits. The -RGAG-Cp recognition motif of one such site, PS1, appears to be selectively protected from X-ray modification in an HBV NCP, compared to a free RNA transcript. These results are consistent with NCP formation *in vitro* occurring via a PS-mediated assembly mechanism. Given that the same RNA and Cp sequences will participate in NCP formation *in vivo*, it seems reasonable to conclude that a similar mechanism contributes to such assembly.

Cryo-EM reconstruction of the JQ707375.1 RNA transcript assembled NCP, reveals extensive density below the globular folds of the Cp shell. Symmetry expansion suggests that there are multiple contacts between this internal density and the outer shell at particle symmetry axes. It is not possible at the resolutions obtained to assign this internal density to either the C-terminal Cp ARD domains, the gRNA transcript or a complex of both. Since this density is not present in NCPs lacking encapsidated RNA^12^, it is clearly a consequence of Cp-gRNA interaction. Previously, we showed that individual oligonucleotides encompassing PS sites will trigger sequence-specific NCP formation *in vitro*. Asymmetric reconstruction of the NCP revealed an inner density larger than could be accounted for by a single RNA stem-loop^12^, although we assumed that PS1, or multiple copies of it, were part of this density. Superposition of that density on the inner layer seen with the gRNA here shows that they are coincident with respect to radial positions, suggesting that this layer does contain RNA. XRF of PS1 confirms this assumption. PSs only need to function, i.e. contact the protein shell of a virion, during assembly. For bacteriophage MS2^48^, many of its RNA PSs dissociate from the CP post-assembly. The fact that PS1 remains in contact after NCP assembly may reflect the need in a para-retrovirus to create an “RNA track” along which the polymerase must move within the NCP as it reverse transcribes the gRNA^12, 50^.

The *in vitro* assembly data with the three variant PS sites suggest that they constitute a novel, evolutionarily stable aspect to NCP assembly regulation. The experiment with the ε variant suggests that gRNA topology is also important in efficient assembly, i.e. that there may be multiple factors that assist the process *in vivo*. The PS-mediated aspect of assembly is a direct anti-viral drug target that could be exploited in the global challenge to treat chronic HBV infections^64^. There are currently only a limited set of clinical interventions, including polymerase inhibitors Lamivudine and Tenofovir, and interferon ^65–67^. Recently novel ligands that target additional aspects of the viral lifecycle have been identified^68–72^. Amongst these are the “capsid assembly modulators, CAMs” that include the hetero-aryl, dihydropyrimidine (HAP) compounds characterised by Zlotnick and colleagues^73, 74^. They have shown that HAPs bind Cp dimers altering their assembly behaviour resulting in formation of “empty” NCP particles. Mathematical modelling suggests that targeting capsid assembly via distinct routes would be synergic^75, 76^. The recent development of a mouse model in which to probe the HBV lifecycle^77^ will hopefully allow these ideas to be tested rapidly allowing the Global Challenge presented by this virus to be met.

## Methods

### Expression and purification of HBV Cp

HBV Cp was expressed in *E. coli* BL21(DE3) cells (T7 Expression strain, New England Biolabs), expressed from a pET28b plasmid. Induction with 1 mM isopropyl-β-D-thiogalactoside (IPTG) at an optical density (OD_600_) of ∼0.6 was followed by growth for 20 h at 21 °C. Cells were lysed using a Soniprep 150 and clarified by spinning at 11,000 xg for 1 h. NCPs were then pelleted by centrifugation at 120,000 xg for 14 h. Pellets were resuspended in 20 mM HEPES (pH 7.5), 250 mM NaCl, 5 mM dithiothreitol (DTT) and applied to a prepacked Captocore 700 column (GE Life Sciences). Fractions containing NCPs were pooled and precipitated with 40% (w/v) ammonium sulfate. NCPs were dissociated into Cp dimers by dialysis into 1.5M guanidinium chloride (GuHCl) as previously described.^12, 38^ All steps after sonication were performed in the presence of complete protease inhibitor tablets (1 tablet per 500 mL buffer, Thermofisher Scientific). Cp dimer concentration was determined by the absorbance at 280 nm (ε_280_ of Cp_2_ = 55,920 L mol^-1^ cm^-1^). Cp fractions with an A_260/280_ ratio of ≤0.65 were used in assembly assays.

### Identification of homologous PS sites in strain JQ707375.1

PS sites within the JQ707375.1, homologous to those found within NC_003977.1, were identified by analysis of stem-loop folds within a 300 nt region of the genome that aligns to the equivalent regions encompassing each PS in the latter strain. A 60 nt reference frame was slid across that region and secondary structure of the corresponding RNA sequence sampled, recording each unique secondary structure fold with negative free energy.^12^ Previously such PS sites were numbered according to the most frequently matched sites in a Bernoulli Plot of the aptamer sequences against the cognate pgRNA.^12^ We retain those labels for the homologous JQ707375.1 strain sites, although this means their numbering does not reflect their relative order. In JQ707375.1, the PS1 equivalent lies between nucleotides 3134-3172 (Fig 2). Mfold suggests that it retains the bulge in the base-paired stem and the -RGAG-loop motif seen in NC_003977.1, although the latter is at the 5′ side of the loop as opposed to being central. This PS1 lies in the gene encoding the X protein, and its nucleotide sequence creates two non-synonymous codons encoding conservative amino acid changes. PS2 (nts 807-859) and PS3 (nts981-1011) sit in regions of the polymerase gene with much lower sequence identities than the overall genome (67 & 75%, respectively). PS2 has a slightly modified loop motif (-RAAG-), which is centrally located, as opposed to being at the 3′ side of the loop, as it is in the NC_003977.1 strain. The loop sits on top of a three base-paired duplex leading to single-stranded bulges of differing sizes. The upper region of this PS stem-loop is homologous to the PS2 oligonucleotide fold in NC_003977.1 (Fig 2). The JQ707375.1 PS3 retains the 3′ positioning of the -RGAG-motif within the loop but has a more extensively base-paired stem than in the previous strain (Fig 2). Nucleotide changes in JQ707375.1 PS2 create one conservative (F to Y), and two semi-conservative (T to N & H to L) amino acid substitutions relative to the NC_003977.1 polymerase. The sequence of the PS3 region yields only synonymous coding changes.

### Preparation of pregenomic RNA

A wild-type gRNA transcript clone was assembled using pAM6 39630^TM^, strain acc. no JQ707375.1, purchased from ATCC®. The RNA sequence was copied from the purchased cDNA in fragments using PCR, and the fragments cloned into the correct order between the *BspHI* and *HindIII* sites in a pACYC184 vector, using a Gibson Assembly® Master Mix, according to the manufacturer’s protocol (New England Biolabs). The sequence of this construct was confirmed by Sanger sequencing (Source Bioscience, Nottingham). HBV pgRNA constructs encompassing PS mutations were produced synthetically, using gene fragments purchased from IDT. Gene fragments were cloned between the *BglII* and *HindIII* sites within an empty pET22b vector using a NEBuilder® HiFi Master Mix, according to the manufacturer’s protocol (New England Biolabs). All pgRNA constructs were designed with a T7 promoter sequence at the 5′ end. Transcription of pgRNA constructs were carried out using a Hiscribe™ T7 High Yield RNA Synthesis Kit (New England Biolabs), after linearization of the DNA plasmid using *HindIII*. RNA was annealed prior to each experiment by heating to 70 °C for 90 secs and cooling slowly to 4 °C in a buffer containing 50 mM NaCl, 10 mM HEPES and 1 mM DTT at pH 7. Products were assessed using a 1% (v/v) denaturing formaldehyde agarose RNA gel. RNA concentration was determined using the A_260_ value (ε_260_ of pgRNA = 32,249,500 L mol^-1^ cm^-1^).

### HBV NCP Assembly Assays

180 μL of 1.1 nM annealed (as above) gRNA transcript in a buffer containing 20 mM HEPES (pH 7.5), 250 mM NaCl and 5 mM DTT, was incubated in each well of a 96-well plate at room temperature for 30 mins. Cp dimer in dissociation buffer (as above) was then titrated into the RNA using a Biomek 4000 liquid handling robot (Beckmann Coulter), step-wise up to a ratio of 1200:1 Cp dimer: RNA. Ten 2 μL titrations were performed, utilising six different stock Cp dimer solutions (indicated in brackets), cumulatively adding 1 nM (100 nM), 10 nM (1 μM), 25 nM (2.5 μM), 75 nM (7.5 μM), 120 nM (12 μM), 240 nM (12 μM), 480 nM (24 μM), 720 nM (24 μM), 960 nM (24 μM) and 1200 nM (24 μM Cp dimer.

This results in a final well volume of 200 μL, a cumulative volume of 19.2 mL/plate, and final concentrations of RNA and Cp of 1 nM and 1.2 μM, respectively. Cp aliquots were calculated to always reach 10% of the final reaction volume limiting the final concentration of GuHCl to 0.15 M in each aliquot. Following incubation at room temperature for 1 hr, the samples were pooled and concentrated to a final volume of 2 mL. The A_260/280_ ratio was measured using a Nanodrop™ One (Thermofisher Scientific) and RNA concentration was calculated using absorbance at 260 nm. This sample was split into 2, with one half treated with 1 µM RNase A, and incubated overnight at 4 °C. After incubation, 5 µL of each sample was visualised by nsEM to assess particle shape and intactness, and the remaining sample analysed by application to a TSK G6000 PWXL column (Tosoh), in a buffer containing 20 mM HEPES (pH 7.5), 250 mM NaCl and 5 mM DTT, using a SEC-MALLS system (ÄKTA Pure (GE Heathcare) connected to a Optilab T-REX refractometer and miniDAWN^®^ multiple angle laser light-scatterer fitted with a Wyatt QELS DLS module (Wyatt Technology)). Light scattering peaks were collected and concentrated to ∼1 ml, where the RNA content of particles and yield considering the RNA input was once again determined using the absorbance at 260 nm. Absorbance values at 260 and 280 were corrected for light-scattering throughout, using the absorbance values at 310 and 340 nm as previously described ^78^.

### R_h_ determination of gRNA transcripts

500 µL of gRNA transcript at a concentration of 500 ng/µL was applied to a TSK G6000 PWXL column (Tosoh) using the buffer and SEC-MALLS system described above.

The hydrodynamic radius of assembled NCPs and gRNA transcripts were determined at the apex of peaks eluted from a TSK G6000 PWXL column. The light scattering at this point is quantified via the second order correlation function, (1) where *I(t)* is the intensity of scattered light at time *t*.

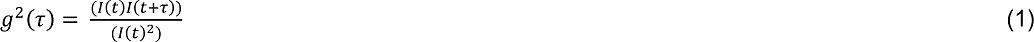

The measured correlation function is fit to equation 2 using a nonlinear least squares fitting algorithm to calculate the decay rate *g*.

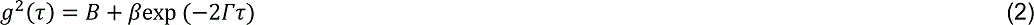

*G* is then converted to the diffusion constant *D* using equation 3, where *q* is the magnitude of the scattering vector given by equation 4.

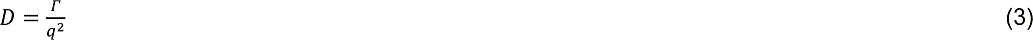

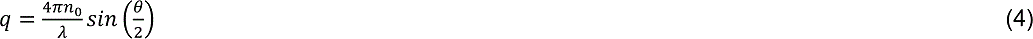

*D* is then fit to the Stokes-Einstein equation (5) to give the R*_h_*.

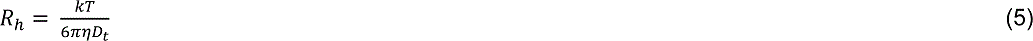

### X-Ray Footprinting of gRNA-containing NCPs

To detect Cp-PS1 interactions in the NCP we used XRF, comparing the footprint on the encapsidated gRNA with the transcript used in the assembly reaction. Both samples were flash frozen and exposed to synchrotron X-rays at the 17-BM beamline (XFP) of the NSLS-II at the Brookhaven National Laboratory (Methods)^26^. X-ray photons are largely absorbed by the solvent water which photolyzes to create hydroxyl radicals that then modify the ribose sugars of solvent accessible nucleotides in a flexibility-dependent manner^25, 26^. Ribose sugars of single-stranded nucleotides are more reactive than those in base-pairs or involved in other molecular contacts. Modification leads to cleavage of the phosphodiester bond^58^, and the frequency of cleavage at each nucleotide can be determined by reverse transcription using fluorescently-tagged primers annealed 3′ to the site of interest.

NCPs were re-assembled as above around the wild-type JQ707375.1 pgRNA in three 96 well plates. These samples were concentrated by centrifugation, and purified over a 10-50% (*w/v*) sucrose density gradient using a SW 40 *Ti* rotor at 190,000 xg for 2 hrs. Purified NCP samples were diluted in 10 mM sodium-phosphate buffer (pH 7.4) to 200 ng/μL with respect to pgRNA using a Nanodrop™ One, and flash-frozen using LN_2_ as 5 μL aliquots in 8 tube PCR strips. Heat-annealed (see above) pgRNA only samples at the same concentration were also flash-frozen as a control.

The samples were footprinted on beamline 17-BM (XFP) at the National Synchrotron Light Source II (Brookhaven National Laboratory, NY, USA). A calibration curve of the X-ray induced photo-bleaching of Alexa488 fluorophore, diluted in 10 mM sodium-phosphate buffer (pH 7.4) was performed to ascertain beam strength, allowing adjustment of sample exposure times between runs to keep levels of RNA cleavage events similar between experiments. Samples were mounted in the beamline in a temperature-controlled (−30 °C, ensuring samples remained frozen), 96-well motorised-holder, which accommodates strips of 8–12 PCR tubes. Beam exposure was controlled using a Uniblitz XRS6 fast shutter (Vincent Associates), exposing samples for 10, 25 and 50 msec, with each time point performed in triplicate. Ideally, exposed samples contained no more than 1 cleavage event across the region of interest, defined as the length of the reverse transcript. This assumes that cleavages elsewhere do not cause large-scale conformational changes in the frozen gRNA within the time span of the exposure (max = 100 milliseconds). Similar data on bacteriophage MS2 gRNA yield predicted secondary structures similar to those seen at atomic resolution by cryoEM^48, 61^, confirming that this is a reasonable assumption.

### Analysis of X-ray induced RNA modification

RNA was extracted from exposed NCPs using the phenol-chloroform (Thermofisher Scientific) extraction technique according to the manufacturers’ protocols, with the exception that the RNAs were precipitated overnight at −20 °C in the presence of 1 volume isopropanol, 0.3 M sodium actetate (pH 5.2) and RNA-grade glycogen (0.01x, Thermofisher Scientific) which gave us higher yields. Recovered RNA was washed 3x with 70%(v/v) ethanol and allowed to air dry for 5 minutes before resuspension with 12 μL nuclease-free water. The beamline-exposed extracted and free pgRNA were reverse transcribed using Superscript IV (Thermofisher Scientific) and a sequence specific 5′ 5(6)-carboxyfluorescein (FAM) labelled primer that attached 3′ of the region of interest. Sequencing ladders were synthesised from *in vitro* transcribed RNA using Hexachloro-fluorescein (HEX) or 5-Carboxytetramethylrhodamine (TAMRA) labelled primers and the addition of a 3:1 molar concentration of ddATP or ddCTP respectively. Bound RNA was degraded with 5 units of RNase H (New England Biolabs) and the cDNA purified by ethanol precipitation overnight at −20 °C (3x volumes of ethanol, 0.3 M sodium acetate 0.01x volume of glycogen). Experimental and sequencing ladder cDNAs were then resuspended in 20 μL formamide, their concentration measured by A_260_ absorbance and 500 ng of each sequencing ladder was spiked into each experimental sample. The samples were heated to 65 °C for 10 minutes then transferred to a 96-well plate and frozen for shipping to DNASeq (Dundee, UK) for capillary electrophoresis (CE).

A reverse primer complementary to pgRNA sites 3328-3308 (sequence 3′ - 5′: AATTTATAAGGGTCAATGTC), was designed to analyse the region encompassing the PS1 signal which was synthesised (IDT) with the appropriate flourophores, (as detailed above) for the aforementioned production of ladders and analysis of beamline-exposed samples.

### CE Analysis

Normalised reactivity profiles were produced from the raw capillary electrophoresis (CE) data using the QuShape Software package^43^ in combination with the BoXFP wrapper module, which is outlined in more detail in Chandler-Bostock *et al.* (unpublished data^41^). For each replicate, pre-processing of the CE data was performed, including signal smoothing, decay correction and alignment of peaks in the ddA sequencing trace to corresponding peaks in the XRF trace. Peaks were identified and peaks in different replicates aligned using the molecular weight size marker. Average and standard deviation of peak intensities were calculated across the replicates to produce the intensity profile. Background correction of samples was performed using the average intensity profile across the unexposed background samples computed. The reactivity profile was calculated and normalised, and error propagation was performed to obtain the error on the normalised intensities. Finally, the ddA traces across all samples from the same primer read was used to generate a consensus sequence of U nucleotide locations across the primer read, which was then aligned to the reference genome to determine the position of the primer read in the genome.

Average Pearson correlation coefficients (PCCs) were calculated across the replicates for each treatment (transcript gRNA = 0.8459278, encapsidated gRNA = 0.94918026), and the normalisation factors used in the computation of the normalised reactivity profiles were recorded for each case (transcript gRNA = 2678.25523, NCP gRNA = 1652.53287). The protection ratio across each primer read was calculated as the normalisation factor of the transcript, divided by the normalisation factor for the encapsidated state of the gRNA, calculated as 1.620697097.

### CryoEM reconstructions of pgRNA containing NCPs

#### Data Acquisition

Lacey carbon 400-mesh copper grids coated with a <3 nm continuous carbon film (Agar Scientific, UK) were glow-discharged in air (10 mA, 30 secs) before applying one 3 μL aliquot of pgRNA containing HBV, reassembled and purified as described above for X-Ray Footprinting. Grids were blotted and vitrified in liquid nitrogen-cooled liquid ethane using a LEICA EM GP plunge freezing device (Leica Microsystems). Chamber conditions were set at 4 °C and 95% relative humidity. Grids were stored in liquid nitrogen prior to imaging with an FEI Titan Krios transmission electron microscope (ABSL, University of Leeds) at 300 kV, at a magnification of 75 000× and a calibrated object sampling of 1.065 Å/pixel. Images were recorded on a FEI Falcon III detector operating in integrating mode. Each movie comprises 59 frames with an exposure rate of 0.9 e^-^ Å^-2^ per frame, with a total exposure time of 1.5 s and an accumulated exposure of 53.1 e^-^ Å^-2^. Data acquisition was performed with EPU Automated Data Acquisition Software for Single Particle Analysis (ThermoFisher) at −0.7 µm to −2.5 µm defocus.

#### Image Processing

The established RELION-3.0 pipeline was used for image processing.^79, 80^ Drift correction was performed using MOTIONCOR2,^81^ and the contrast transfer function estimated using Gctf.^82^ A subset of 1000 particles were picked and classified using reference-free 2D classification and used as templates for the RELION autopicking procedure. Picked particles were sorted using 2D classification, with 127,410 *T* =4 and 23,257 *T=*3 particles subsequently extracted. The large majority (∼15,000) of *T*=3 particles were discarded due to heterogeneity. 104,337 *T*=4 particles were selected and subjected to 3D classification and subsequent auto-refinement with icosahedral symmetry (I1) imposed and without (C1), using EMD-3715 low pass filtered to 60 Å resolution as a reference model. This reconstruction was post-processed to mask and correct for the B-factor of the map. The CTF refinement routine implemented in RELION-3.0^79^ was used to refine the reconstruction further, yielding a map with an overall resolution at 3.2 Å, based on the gold-standard (FSC = 0.143) criterion.

To investigate the density of the pgRNA, a focussed 3D classification approach was employed. Each particle contributing to the final icosahedral symmetry-imposed reconstruction was assigned 60 orientations corresponding to its icosahedrally-related views using the relion_symmetry_expand tool. SPIDER^48^ was used to generate a spherical mask placed just beneath the Cp shell of the NC, and the symmetry expanded particles were subjected to masked 3D classification, sorting them into 5 classes without alignment, using a regularisation parameter of 25. Particles from these classes were reconstructed using the relion_reconstruct tool, without imposing symmetry, and postprocessed, yielding maps with a resolution of 3.5 Å. UCSF Chimera was used for visualisation and figure generation.^83^

#### Model building and refinement

The structure of HBV NC (PDB 3J2V)^54^ was first manually docked as a rigid body into the density and followed by real space fitting with the Fit in Map routine in UCSF Chimera.^83^ A first step of real space refinement was performed in Phenix.^84^ The model was then manually rebuilt in Coot^85^ to optimize the fit to the density. After icosahedral symmetrisation to generate the entire capsid, a second step of real space refinement was performed in Phenix. Refinement statistics are listed in Supplementary Table 2.

#### Model validation and analysis

The FSC curve between the final model and map after post-processing in RELION (Model vs Map), is shown in Sup Fig 3B. To perform cross-validation against overfitting, the atoms in the final atomic model were displaced by 0.5 Å in random directions using Phenix. The shifted coordinates were then refined against one of the half-maps (work set) in Phenix using the same procedure as for the refinement of the final model. The other half-map (test set) was not used in refinement for cross-validation. FSC curves of the refined shifted model against the work set (FSCwork) and against the test set (FSCtest), are shown in Sup Fig 3B. The FSCwork and FSCtest curves are not significantly different, consistent with the absence of overfitting in the final models. The quality of the atomic model, including basic protein geometry, Ramachandran plots, clash analysis, was assessed and validated with Coot, MolProbity^86^ as implemented in Phenix, and with the Worldwide PDB (wwPDB) OneDep System (https://deposit-pdbe.wwpdb.org/deposition). Graphics were produced by UCSF Chimera.^83^

## Data availability

The data that supports the findings of this study are available from the corresponding authors upon request. Correspondence and requests for materials should be addressed to N.P or P.G.S. The atomic coordinates for HBV NC were deposited in the Protein Data Bank with code 7ABL. The icosahedrally averaged, symmetry-expanded and focused classified on genome and asymmetric cryo-EM density maps were deposited in the EM Data Bank with codes EMD-11700, EMD-11702 and EMD-11701, respectively.

## Author contributions

EUW and SC analysed XRF data, CPM helped analyse cryo-EM data and produced structural figures. EF and JB oversaw experiments at the Brookhaven National Laboratory. NP performed experiments and analysed data. NP and PGS wrote the paper with help from RT, NAR and CPM. NP, PGS and RT conceived the project.

## Acknowledgment

We thank Prof Adam Zlotnick, Indiana University, for the gift of his Cp expression construct and advice on its purification, and Ms Leah Wells who assisted with reassembly experiments during her BSc (Honours) undergraduate research project. We thank the Medical Research Foundation for the award of a career development grant to NP, and the UK MRC for previous grant funding to study HBV assembly (MRF-044-0002-RG-PATEL & MR/N021517/1). RT & PGS thank The Wellcome Trust (Joint Investigator Award Nos. 110145 & 110146 to PGS & RT, respectively) for funding, and we acknowledge the financial support of The Trust of infrastructure and equipment in the Astbury Centre, University of Leeds (089311/Z/09/Z; 090932/Z/09/Z & 106692), and for their additional support, together with The University of Leeds, of the Astbury Biostructure Facility. RT acknowledges additional funding via an EPSRC Established Career Fellowship (EP/R023204/1) and a Royal Society Wolfson Fellowship (RSWF\R1\180009). Portions of this work used the XFP (17-BM) beamline at NSLS-II. Development of XFP was made possible by the National Science Foundation, Division of Biological Infrastructure (grant No. 1228549), while operations support of XFP was provided by the National Institutes of Health (grant No. P30-EB-009998). NSLS-II, a US Department of Energy (DOE) Office of Science User Facility operated for the DOE Office of Science by Brookhaven National Laboratory, was supported under Contract No. DE-SC0012704. We thank DNA Sequencing & Services (MRC I PPU, School of Life Sciences, University of Dundee, Scotland, www.dnaseq.co.uk) for DNA sequencing.

## Supplementary Information

### Supplementary Methods

#### Motif analysis in other packaged sequences

Three sequences were considered in this analysis: the LacZ sequence which has been shown *in vivo* not to package into HBV virions ^1^; the LacZ sequence, with a 5’ HBV genomic fragment containing ε which does get packaged into HBV virions^1^; and a minimal sequence shown to package into HBV virions^2^. Each of these sequences was searched for occurrences of the RGAG and GAAG Cp-binding motifs seen in the PS motifs of strain JQ707375.1. The potential secondary structures within these sequences were determined using Sfold^3^, both globally across the entire sequence and locally using a sliding window of 80 nts. For both global folds and each 80 nt window 1000 sample folds were calculated. The frequencies of folds that presenting the Cp-recognition motifs in a loop was then recorded.

Sequences tested-

### Epsilon (red)-LacZ sequence

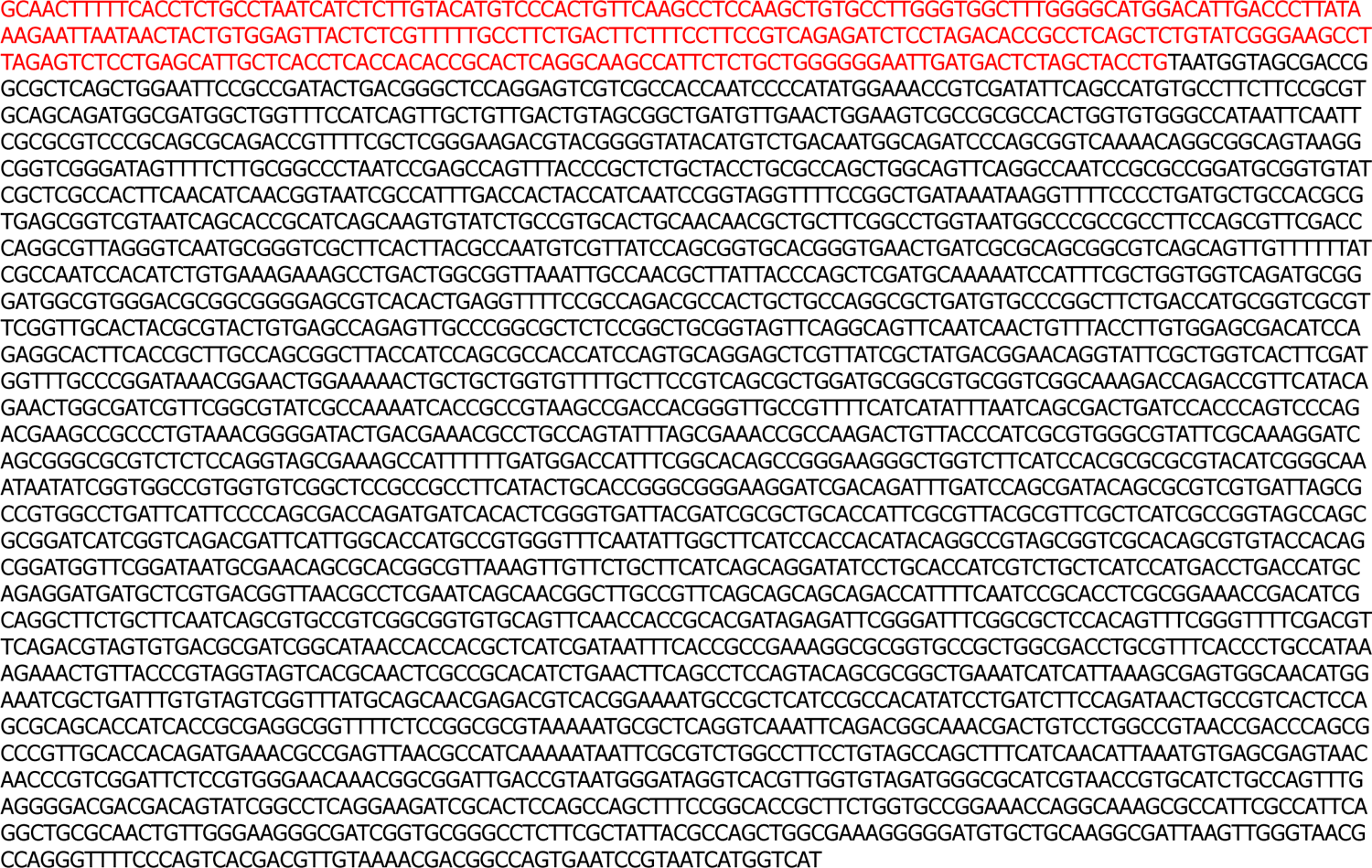

### LacZ sequence

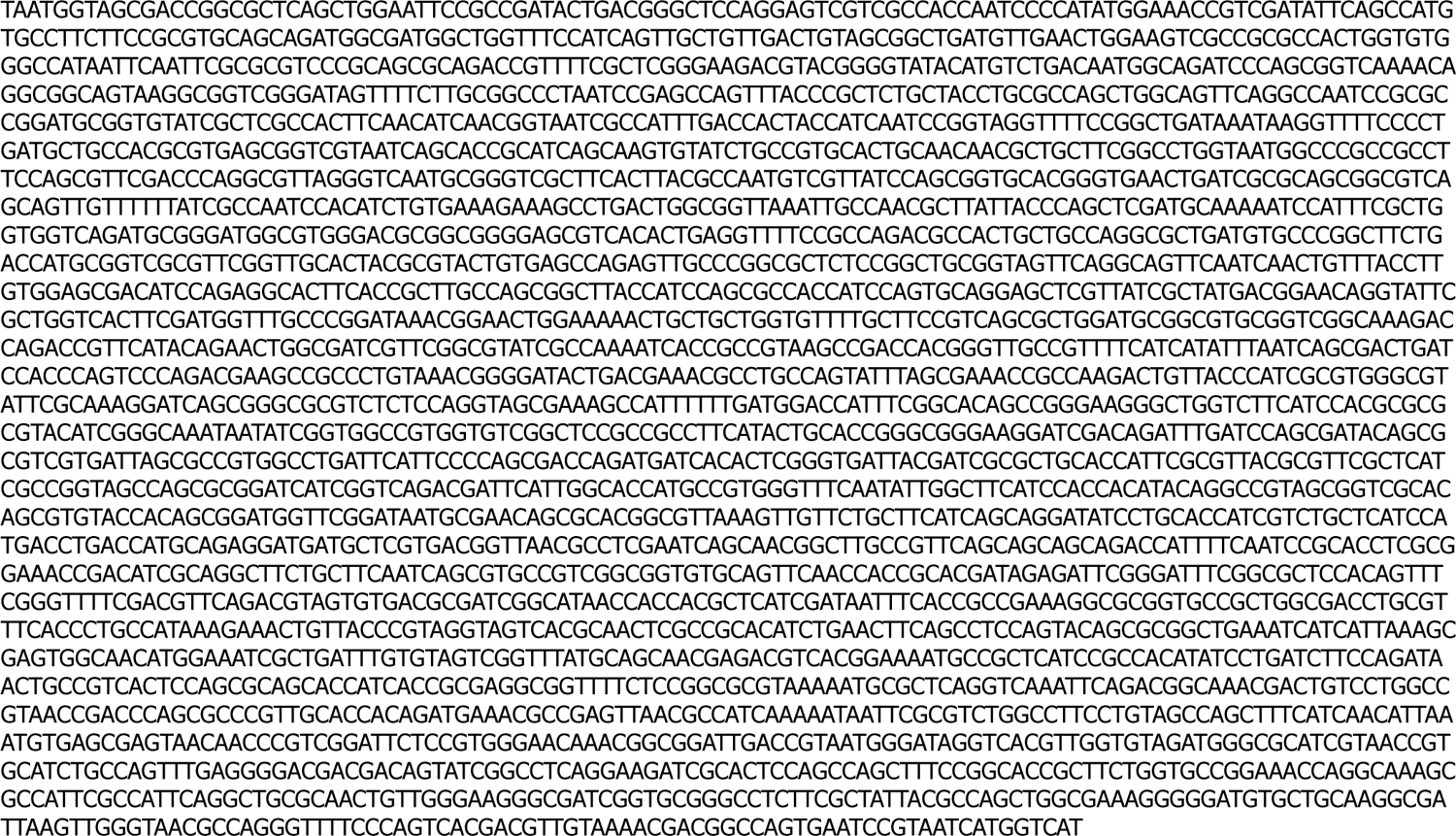

### Minimal Junker sequence^2^

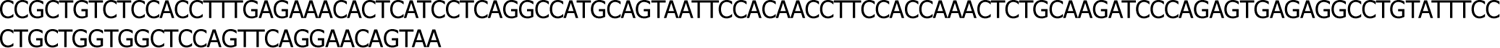

### Supplementary Figures

**Supplementary Figure 1:**
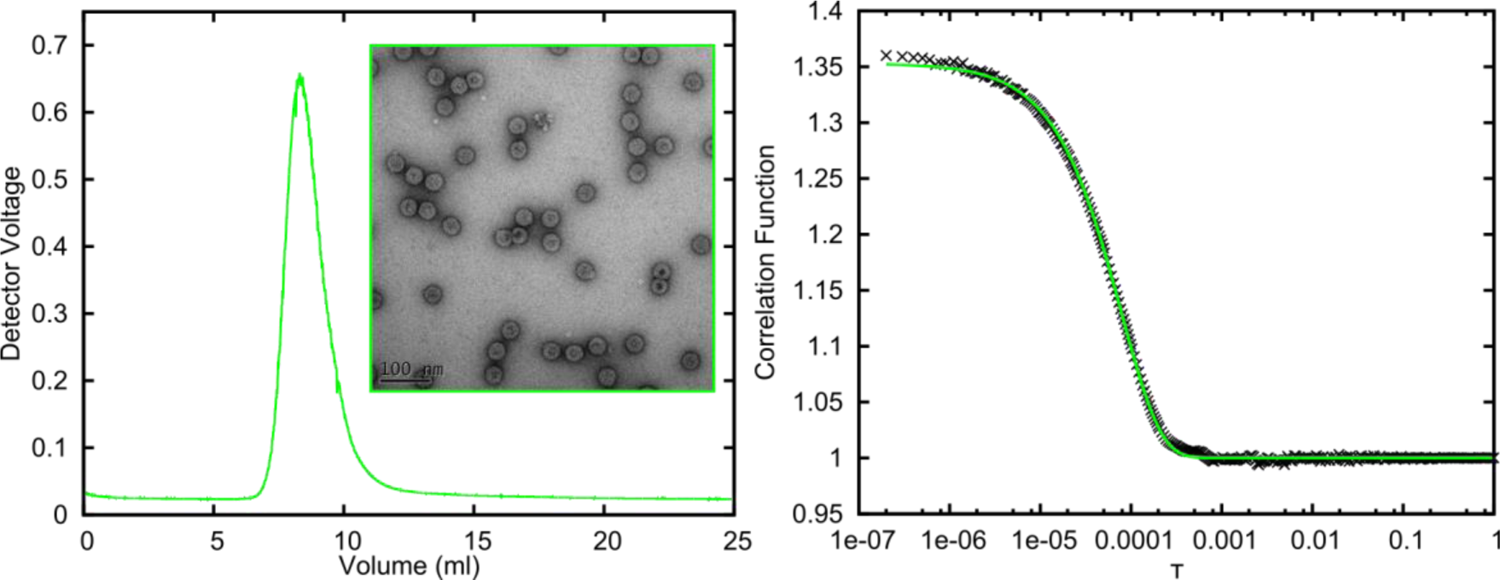
Light-scattering, gel filtration signals from HBV NCPs formed in *E. coli*. LS and R*_h_* data from SEC-MALLS analysis of HBV NCPs formed up on Cp expression in *E. coli* (R*_h_* = 19.4 nm).

**Supplementary Figure 2:**
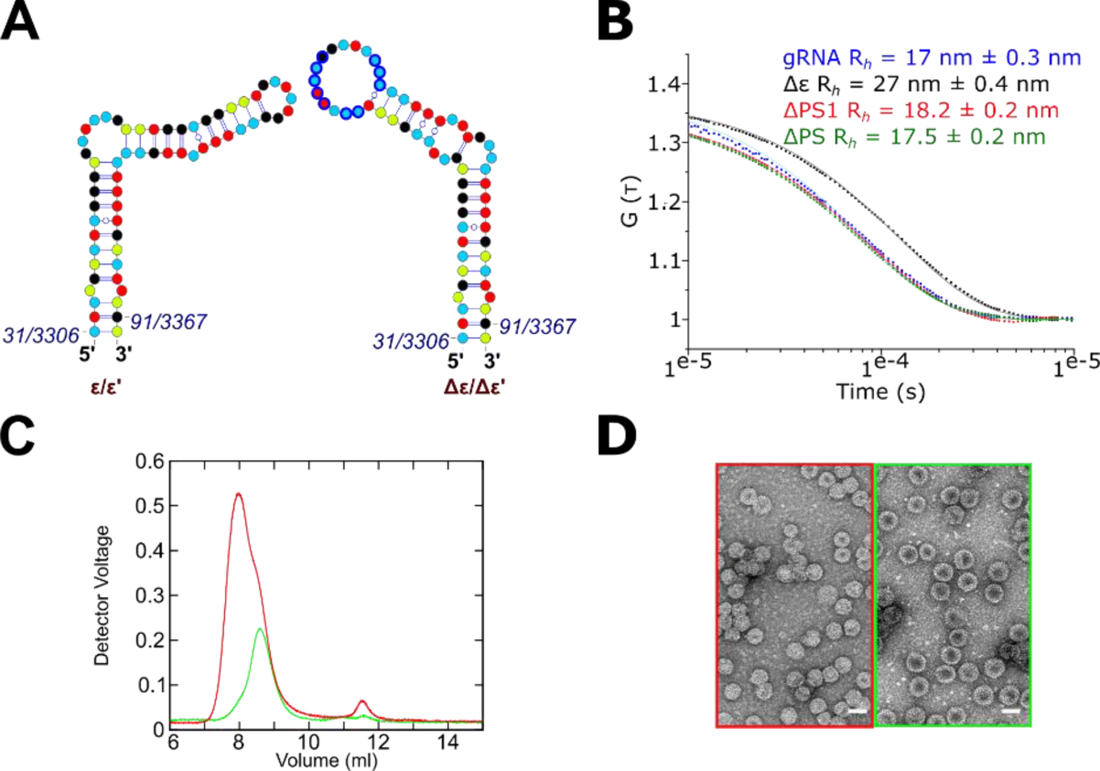
Reassembly of NCPs with Δɛ variant pgRNA. S-fold secondary structures of the regions surrounding: (A) the wild-type JQ707375.1 ɛ/ ɛ′ *(left)* and the Δɛ variant *(right)*. RNA nucleotides are shown as in Fig 2. (B) Autocorrelation curves for the JQ707375.1 transcript (*blue*), and the ΔPS (*green*), ΔPS1 (*red*) and Δɛ (*black*) variants measured by SEC-MALLs. The derived R*_h_* values for these RNAs: gRNA transcript, ΔPS, ΔPS1 and Δɛ, are 17 ± 0.3, 17.5 ± 0.2, 18.2 ± 0.2 and 27± 0.4 nm, respectively. (C) Result of an *in vitro* reassembly of the Δɛ variant, as described for the gRNA transcript (Fig. 3A). (D) *Left / Right:* Colour-coded nsEMs of products from (C). Scale bars = 50 nm.

**Supplementary Figure 3:**
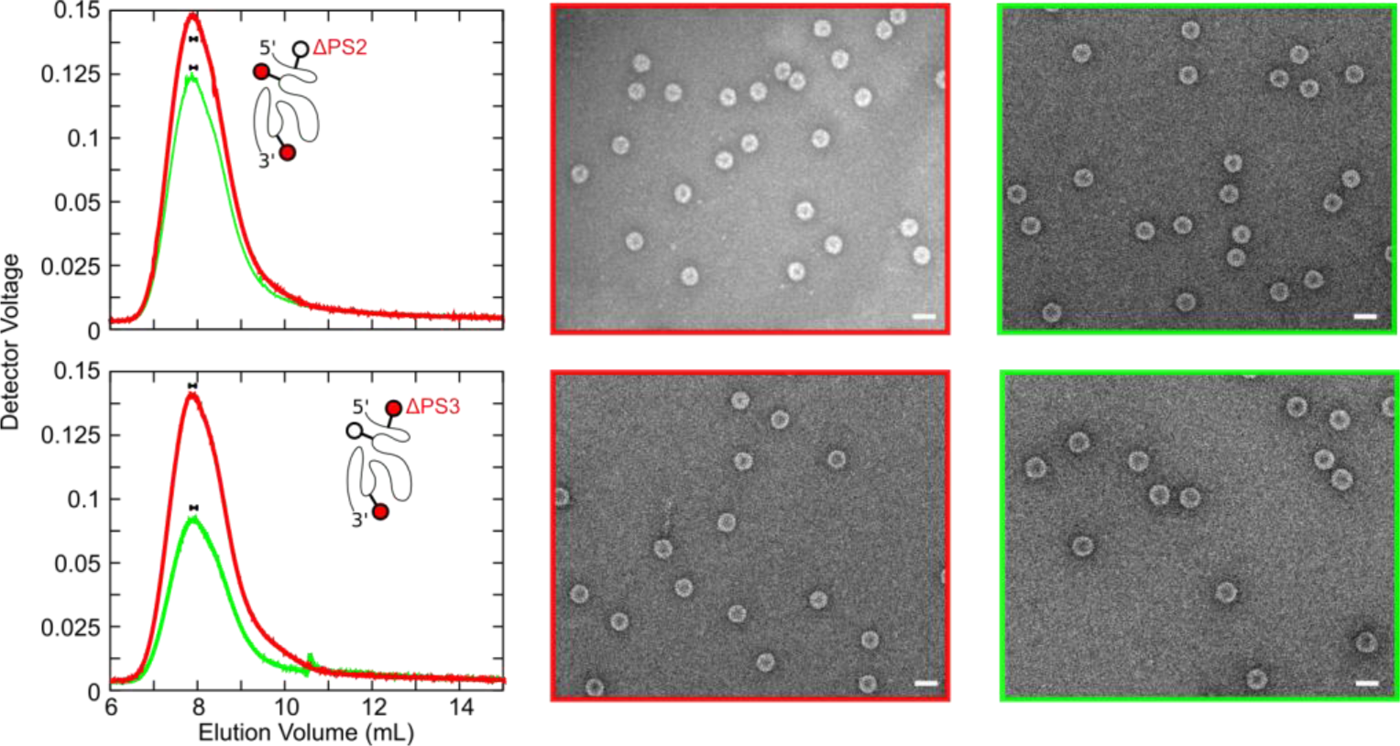
Reassembly of NCPs with pgRNAs containing variants of PS2 or PS3. NCP reassemblies containing 1 nM pgRNA with ΔPS2 (top) or ΔPS3 (bottom) and HBV Cp dimer titrations, see Fig. 2A for details. LS traces before (red) or after (green) 1 µM RNase A treatment. *Middle / Right panels:* nsEMs of re-assembly products, colour-coded as in LS traces. Scale bars = 50 nm.

**Supplementary Figure 4:**
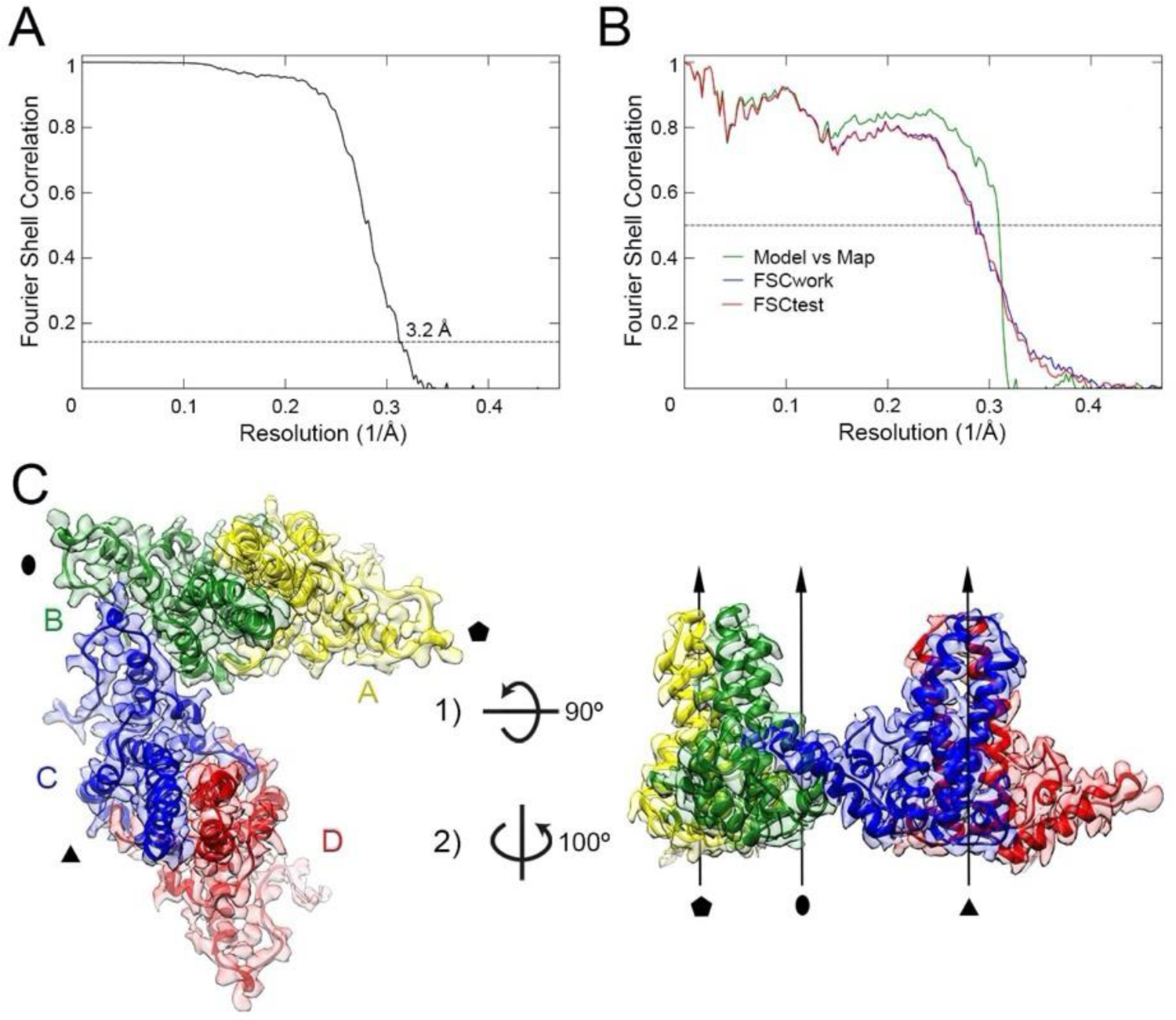
Resolution and model validation of HBV *T*=4 NCP structure. (A) Fourier Shell Correlation (FSC) resolution curve for the icosahedrally-averaged 3D reconstruction of HBV *T*=4 NCP. Resolution based on the gold standard 0.143 criterion is 3.2 Å. (B) Cross-validation against overfitting of the model. The FSC curve for the final atomic model refined against the post-processed map (green curve, Model vs Map), and FSC curves for the randomly shifted and refined atomic model against the half map used in the refinement (blue curve, FSCwork) and against the half map not used in the refinement (red curve, FSCtest). (C) Atomic model of the asymmetric unit of HBV *T*=4 NCP shown as ribbon diagrams (top view, left; side view, right) colour-coded as in Fig. 5, fitted into the 3.2 Å resolution cryo-EM density map shown as colour-coded semi-transparent surface. Symbols and arrows indicate icosahedral symmetry axes.

**Supplementary Figure 5:**
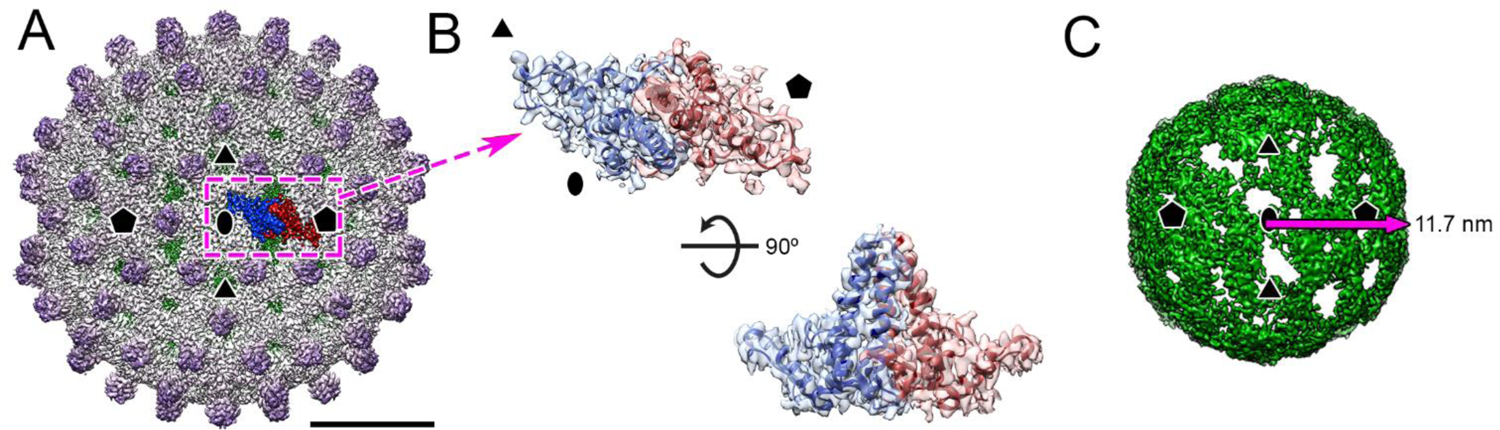
Cryo-EM reconstruction of the *T*=4 NCP formed with the gRNA transcript. (A) Front-half of the icosahedrally-averaged cryo-EM density map of the reassembled HBV *T*=4 NCP containing the wild-type JQ707375.1 RNA transcript at 3.2 Å resolution (bar = 100 Å). The dashed box surrounds a Cp dimer, subunits highlighted in blue and red, shown in (B) with top and side views of the dimer (PDB: 7ABL) fitted into the map segmented from (A). (C) One of the selected class (Class 3) obtained after symmetry expansion and focused classification of the internal density of the structure in (A) low-pass filtered to 5 Å resolution. The arrow indicates the radius of this feature. Maps are radially colour-coded (green-white-purple), shown at 2σ, and viewed along a two-fold axis.

**Supplementary Figure 6:**
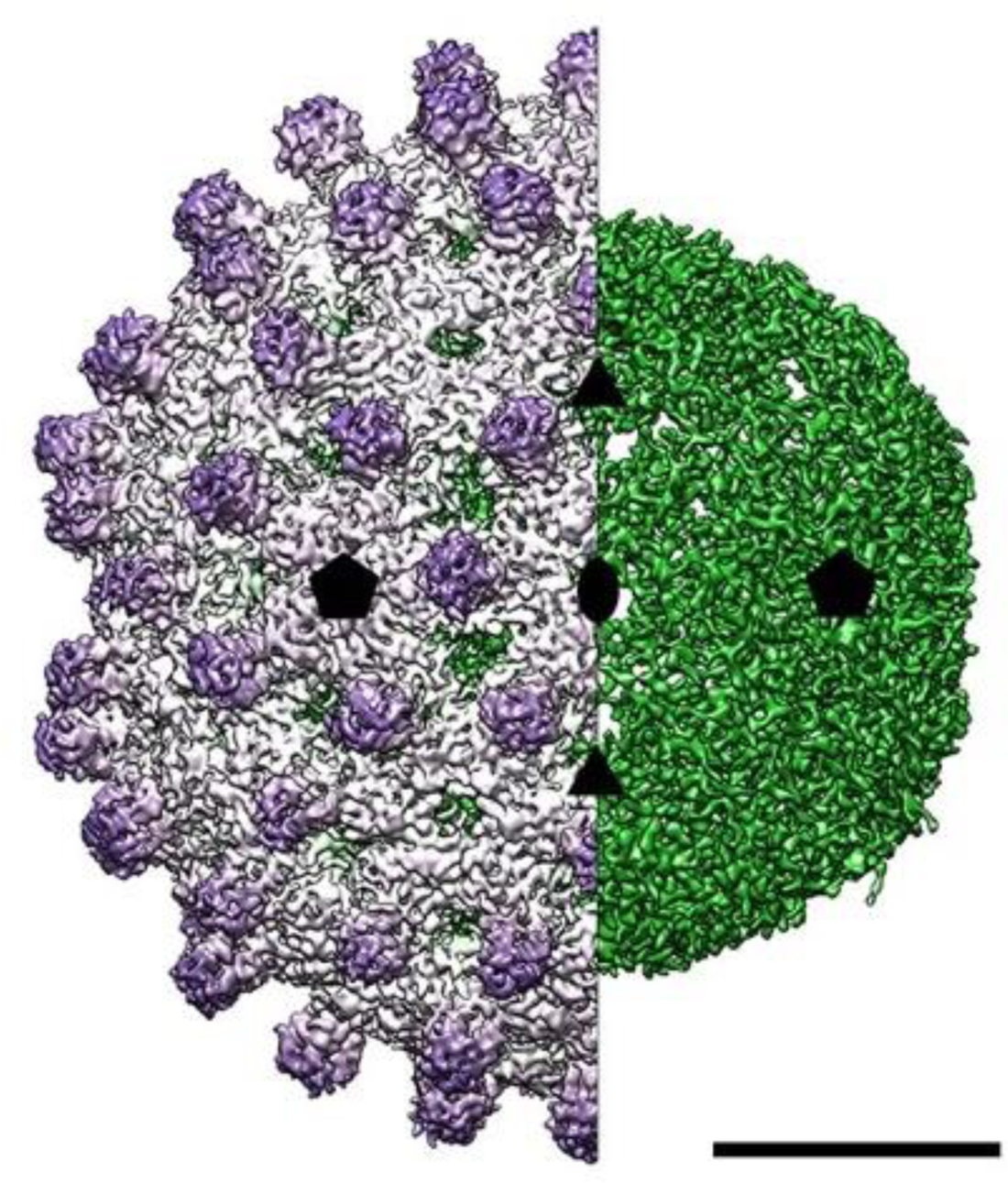
Asymmetric reconstruction of the *T*=4 NCP. Asymmetric cryo-EM density map of HBV *T*=4 NCP reconstructed at 4.2 Å resolution without imposition of icosahedral symmetry (right half, internal density in Sup Fig 5 low-pass filtered to 5 Å resolution). Bar = 100 Å.

**Supplementary Figure 7:**
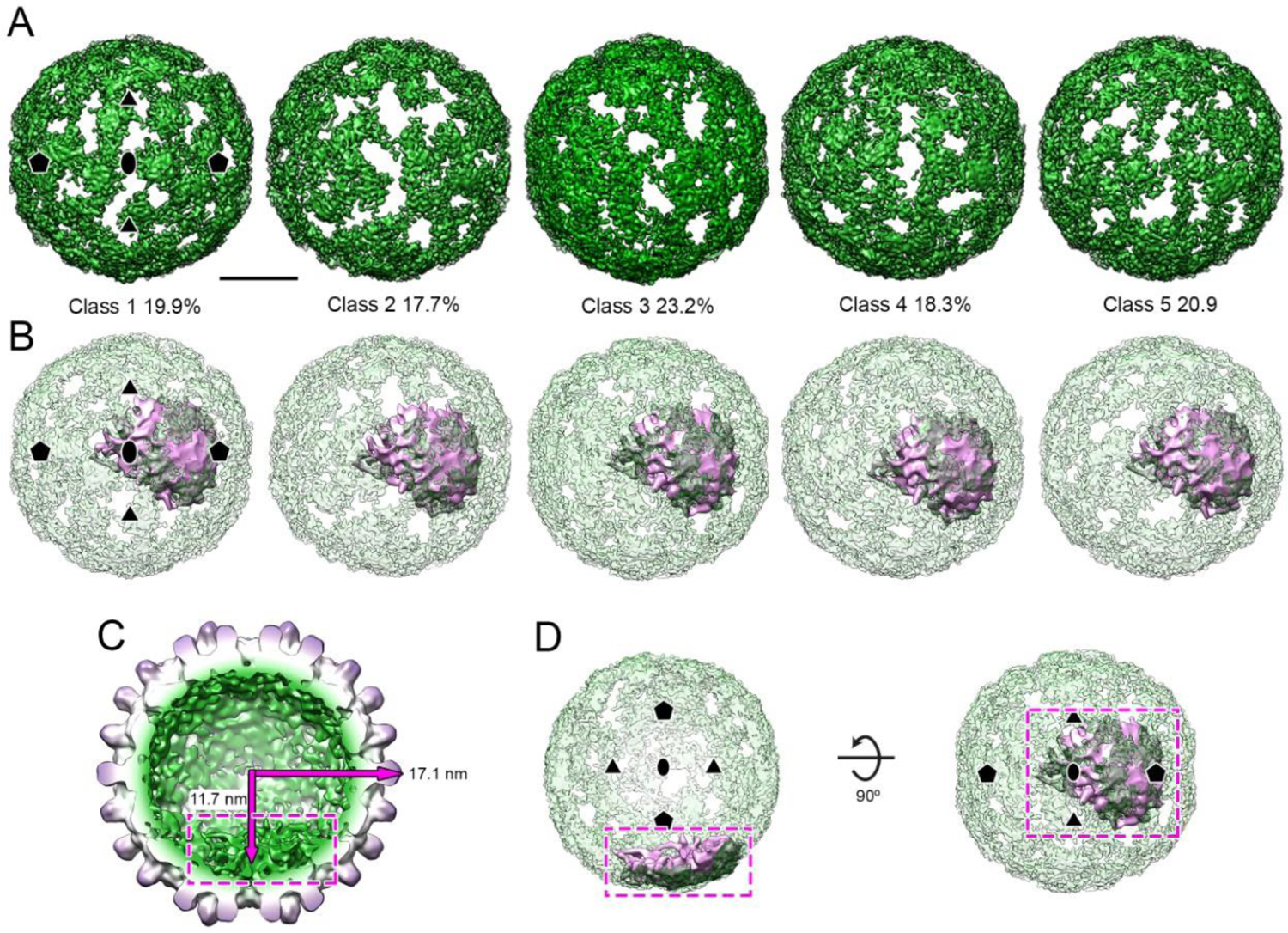
Evidence that the internal density may include the RNA PSs. (A) Symmetry expansion and focused classification of the internal density (right half, low-pass filtered to 5 Å resolution) of the structure in Figure 4. All particles fit in one of five, equally-populated similar classes, shown below. Bar = 100 Å. Symbols indicate icosahedral symmetry axes. (B) Superposition of the density corresponding to the asymmetric feature seen in NC_003977.1 NCPs assembled around an oligonucleotide encompassing the PS1 from that strain (pink) into the internal density (transparent green) from classes 1 to 5^4^. (C) Back-half of the asymmetric cryo-EM density map of PS1 containing HBV VLP at 11.4 Å resolution (EMD-3714). Dashed box indicates the density corresponding to the feature observed in the NCP formed around PS1 (B). Arrows indicate the radii of the NCP and the internal shell. (D) Detail of the superposition of density corresponding to the PS1 NCP (pink) into internal shell (transparent green) of density from Class 3. Maps are radially colour-coded as in Fig.4, shown at 2σ and viewed along a two-fold axis.

**Supplementary Table 1:** RGAG and GAAG motifs found in Epsilon+LacZ. Left to right: position of motif in the sequence; motif sequence; region in which the motif is located; number of times this motif is expressed in a loop when the entire sequence is folded with Sfold; number of times this motif is expressed in a loop when the sequence is folded locally with Sfold using a sliding window of 80 nts.

| Motif position | Motif sequence | Motif Region | Loop expression frequency |  |
| --- | --- | --- | --- | --- |
|  |  |  | Global folding | Local folding |
| 124 | GGAG | Epsilon | 1 | 15 |
| 167 | AGAG | Epsilon | 0 | 76 |
| 203 | GAAG | Epsilon | 76 | 1732 |
| 211 | AGAG | Epsilon | 3 | 104 |
| 351 | GGAG | LacZ | 0 | 0 |
| 480 | GAAG | LacZ | 0 | 10 |
| 552 | GAAG | LacZ | 0 | 289 |
| 1154 | GGAG | LacZ | 0 | 48 |
| 1262 | AGAG | LacZ | 4 | 6 |
| 1324 | GGAG | LacZ | 0 | 14 |
| 1336 | AGAG | LacZ | 0 | 32 |
| 1391 | GGAG | LacZ | 1 | 0 |
| 1647 | GAAG | LacZ | 2 | 100 |
| 1809 | GAAG | LacZ | 59 | 2958 |
| 1903 | GAAG | LacZ | 0 | 10 |
| 2260 | AGAG | LacZ | 0 | 5 |
| 2420 | AGAG | LacZ | 3 | 496 |
| 3110 | GAAG | LacZ | 0 | 11 |
| 3202 | GAAG | LacZ | 0 | 1556 |

**Supplementary Table 2:** RGAG and GAAG motifs found in LacZ. Left to right: position of motif in the sequence, motif sequence, number of times this motif is expressed in a loop when the entire sequence is folded with Sfold, number of times this motif is expressed in a loop when the sequence is folded locally with Sfold using a sliding window of 80 nts.

| Motif position | Motif sequence | Loop expression frequency |  |
| --- | --- | --- | --- |
|  |  | Global folding | Local folding |
| 54 | GGAG | 0 | 2 |
| 183 | GAAG | 0 | 10 |
| 255 | GAAG | 0 | 245 |
| 857 | GGAG | 0 | 86 |
| 965 | AGAG | 0 | 9 |
| 1027 | GGAG | 0 | 3 |
| 1039 | AGAG | 0 | 31 |
| 1094 | GGAG | 0 | 2 |
| 1350 | GAAG | 1 | 57 |
| 1512 | GAAG | 63 | 2404 |
| 1606 | GAAG | 0 | 9 |
| 1963 | AGAG | 0 | 4 |
| 2123 | AGAG | 1 | 482 |
| 2813 | GAAG | 0 | 5 |
| 2905 | GAAG | 0 | 1147 |

**Supplementary Table 3:** RGAG motifs found in Junker minimal sequence. Left to right: position of motif in the sequence, motif sequence, number of times this motif is expressed in a loop when the entire sequence is folded with Sfold, number of times this motif is expressed in a loop when the sequence is folded locally with Sfold using a sliding window of 80 nts.

| Motif position | Motif sequence | Loop expression frequency |  |
| --- | --- | --- | --- |
|  |  | Global folding | Local folding |
| 83 | AGAG | 0 | 79 |
| 89 | AGAG | 0 | 124 |

## References

1. World Health Organization. Weekly epidemiological record. Hepatitis B Vaccines. 84, 405–420 (2009).

2. Tillmann, H. L. Antiviral therapy and resistance with hepatitis B virus infection. World J. Gastroenterol. (2007) doi:10.3748/wjg.v13.i1.125.

3. Murray, K. et al. Protective immunisation against hepatitis B with an internal antigen of the virus. J. Med. Virol. (1987) doi:10.1002/jmv.1890230202.

4. World Health Organization. Progress report on HIV, viral hepatitis and sexually transmitted infections, 2019. **1**, 1–39 (2019).

5. Nassal, M. Hepatitis B viruses: Reverse transcription a different way. Virus Res. (2008) doi:10.1016/j.virusres.2007.12.024.

6. Lok, A. S. et al. Long-term safety of lamivudine treatment in patients with chronic hepatitis B. Gastroenterology 125, 1714–1722 (2003).

7. World Health Organization. Combating hepatitis B and C to reach elimination by 2030. 1–16 (2016).

8. Bock, C. T. et al. Structural organization of the hepatitis B virus minichromosome. J Mol Biol 307, 183–196 (2001).

9. Guo, Y. H., Li, Y. N., Zhao, J. R., Zhang, J. & Yan, Z. HBc binds to the CpG islands of HBV cccDNA and promotes an epigenetic permissive state. Epigenetics 6, 720–726 (2011).

10. Selzer, L. & Zlotnick, A. Assembly and Release of Hepatitis B Virus. Cold Spring Harb Perspect Med 5, (2015).

11. Seeger, C. & Mason, W. S. Hepatitis B virus biology. Microbiol Mol Biol Rev 64, 51–68 (2000).

12. Patel, N. et al. HBV RNA pre-genome encodes specific motifs that mediate interactions with the viral core protein that promote nucleocapsid assembly. Nat. Microbiol. 2, 17098 (2017).

13. Bunka, D. H. J. & Stockley, P. G. Aptamers come of age - At last. Nature Reviews Microbiology (2006) doi:10.1038/nrmicro1458.

14. Stockley, P. G. et al. Packaging signals in single-stranded RNA viruses: nature’s alternative to a purely electrostatic assembly mechanism. J Biol Phys 39, 277–287 (2013).

15. Dykeman, E. C., Stockley, P. G. & Twarock, R. Packaging signals in two single-stranded RNA viruses imply a conserved assembly mechanism and geometry of the packaged genome. J Mol Biol 425, 3235–3249 (2013).

16. Borodavka, A., Tuma, R. & Stockley, P. G. A two-stage mechanism of viral RNA compaction revealed by single molecule fluorescence. RNA Biol 10, 481–489 (2013).

17. Bunka, D. H. et al. Degenerate RNA packaging signals in the genome of Satellite Tobacco Necrosis Virus: implications for the assembly of a T=1 capsid. J Mol Biol 413, 51–65 (2011).

18. Rolfsson, O. et al. Direct Evidence for Packaging Signal-Mediated Assembly of Bacteriophage MS2. J Mol Biol (2015) doi:10.1016/j.jmb.2015.11.014.

19. Shakeel, S. et al. Genomic RNA folding mediates assembly of human parechovirus. Nat Commun 8, 5 (2017).

20. R, T. & PG, S. RNA-Mediated Virus Assembly: Mechanisms and Consequences for Viral Evolution and Therapy. Annu. Rev. Biophys. 48, 495–514 (2019).

21. Conley, M. J. et al. Calicivirus VP2 forms a portal-like assembly following receptor engagement. Nature (2019) doi:10.1038/s41586-018-0852-1.

22. Scheres, S. H. W. Processing of Structurally Heterogeneous Cryo-EM Data in RELION. In Methods in Enzymology (2016). doi:10.1016/bs.mie.2016.04.012.

23. McElwee, M., Vijayakrishnan, S., Rixon, F. & Bhella, D. Structure of the herpes simplex virus portal-vertex. PLoS Biol. (2018) doi:10.1371/journal.pbio.2006191.

24. Zhou, M. et al. Atomic structure of the apoptosome: Mechanism of cytochrome c- and dATP-mediated activation of Apaf-1. Genes Dev. (2015) doi:10.1101/gad.272278.115.

25. Adilakshmi, T., Soper, S. F. C. & Woodson, S. A. Biophysical, Chemical, and Functional Probes of RNA Structure, Interactions and Folding: Part A. (Elsevier, 2009).

26. Asuru, A. et al. The XFP (17-BM) beamline for X-ray footprinting at NSLS-II. J. Synchrotron Radiat. (2019) doi:10.1107/S1600577519003576.

27. Tetter, S. et al. Evolution of a virus-like architecture and packaging mechanism in a repurposed bacterial protein. Science (80-.). 372, 1220–1224 (2021).

28. Bartenschlager, R. & Schaller, H. Hepadnaviral assembly is initiated by polymerase binding to the encapsidation signal in the viral RNA genome. EMBO J 11, 3413–3420 (1992).

29. Gazina, E. V., Fielding, J. E., Lin, B. & Anderson, D. A. Core Protein Phosphorylation Modulates Pregenomic RNA Encapsidation to Different Extents in Human and Duck Hepatitis B Viruses. J. Virol. (2000) doi:10.1128/jvi.74.10.4721-4728.2000.

30. Lan, Y. T., Li, J., Liao, W. & Ou, J. Roles of the three major phosphorylation sites of hepatitis B virus core protein in viral replication. Virology 259, 342–348 (1999).

31. Junker-Niepmann, M., Bartenschlager, R. & Schaller, H. A short cis-acting sequence is required for hepatitis B virus pregenome encapsidation and sufficient for packaging of foreign RNA. EMBO J. (1990) doi:10.1002/j.1460-2075.1990.tb07540.x.

32. Pollack, J. R. & Ganem, D. An RNA stem-loop structure directs hepatitis B virus genomic RNA encapsidation. J. Virol. (1993) doi:10.1128/jvi.67.6.3254-3263.1993.

33. M. Zuker. Mfold web server for nucleic acid folding and hybridization prediction. Nucleic Acids Res. (2003).

34. Thai, H. et al. Convergence and coevolution of Hepatitis B virus drug resistance. Nat. Commun. 3, 789 (2012).

35. Patel, N. et al. Revealing the density of encoded functions in a viral RNA. Proc Natl Acad Sci U S A 112, 2227–2232 (2015).

36. Routh, A., Domitrovic, T. & Johnson, J. E. Host RNAs, including transposons, are encapsidated by a eukaryotic single-stranded RNA virus. Proc Natl Acad Sci U S A 109, 1907–1912 (2012).

37. Ford, R. J. et al. Sequence-specific, RNA-protein interactions overcome electrostatic barriers preventing assembly of satellite tobacco necrosis virus coat protein. J Mol Biol 425, 1050–1064 (2013).

38. Porterfield, J. Z. et al. Full-length hepatitis B virus core protein packages viral and heterologous RNA with similarly high levels of cooperativity. J Virol 84, 7174–7184 (2010).

39. Borodavka, A., Tuma, R. & Stockley, P. G. Evidence that viral RNAs have evolved for efficient, two-stage packaging. Proc Natl Acad Sci U S A 109, 15769–15774 (2012).

40. Garmann, R. F. et al. Role of electrostatics in the assembly pathway of a single-stranded RNA virus. J Virol 88, 10472–10479 (2014).

41. Rudnick, J. & Bruinsma, R. Icosahedral packing of RNA viral genomes. Phys. Rev. Lett. (2005) doi:10.1103/PhysRevLett.94.038101.

42. van der Schoot, P. & Bruinsma, R. Electrostatics and the assembly of an RNA virus. Phys Rev E Stat Nonlin Soft Matter Phys 71, 61928 (2005).

43. Belyi, V. A. & Muthukumar, M. Electrostatic origin of the genome packing in viruses. Proc. Natl. Acad. Sci. U. S. A. (2006) doi:10.1073/pnas.0608311103.

44. Balint, R. & Cohen, S. S. The incorporation of radiolabeled polyamines and methionine into turnip yellow mosaic virus in protoplasts from infected plants. Virology 144, 181–193 (1985).

45. Bruinsma, R. F. Physics of RNA and viral assembly. Eur Phys J E Soft Matter 19, 303–310 (2006).

46. Twarock, R. & Stockley, P. G. RNA-Mediated Virus Assembly: Mechanisms and Consequences for Viral Evolution and Therapy. Annual Review of Biophysics (2019) doi:10.1146/annurev-biophys-052118-115611.

47. Dykeman, E. C., Stockley, P. G. & Twarock, R. Solving a Levinthal’s paradox for virus assembly identifies a unique antiviral strategy. Proc Natl Acad Sci U S A 111, 5361–5366 (2014).

48. Dai, X. et al. In situ structures of the genome and genome-delivery apparatus in a single-stranded RNA virus. Nature 541, 112–116 (2017).

49. Meng, R. et al. Structural basis for the adsorption of a single-stranded RNA bacteriophage. Nat. Commun. 10, 1–8 (2019).

50. Wang, J. C., Nickens, D. G., Lentz, T. B., Loeb, D. D. & Zlotnick, A. Encapsidated hepatitis B virus reverse transcriptase is poised on an ordered RNA lattice. Proc Natl Acad Sci U S A 111, 11329–11334 (2014).

51. Wynne, S. A., Crowther, R. A. & Leslie, A. G. W. The crystal structure of the human hepatitis B virus capsid. Mol. Cell (1999) doi:10.1016/S1097-2765(01)80009-5.

52. Böttcher, B., Wynne, S. A. & Crowther, R. A. Determination of the fold of the core protein of hepatitis B virus by electron cryomicroscopy. Nature (1997) doi:10.1038/386088a0.

53. Conway, J. F. et al. Visualization of a 4-helix bundle in the hepatitis B virus capsid by cryo-electron microscopy. Nature (1997) doi:10.1038/386091a0.

54. Yu, X., Jin, L., Jih, J., Shih, C. & Hong Zhou, Z. 3.5Å cryoEM Structure of Hepatitis B Virus Core Assembled from Full-Length Core Protein. PLoS One (2013) doi:10.1371/journal.pone.0069729.

55. Conley, M. J. & Bhella, D. Asymmetric analysis reveals novel virus capsid features. Biophysical Reviews (2019) doi:10.1007/s12551-019-00572-9.

56. Ruan, L., Hadden, J. A. & Zlotnick, A. Assembly Properties of Hepatitis B Virus Core Protein Mutants Correlate with Their Resistance to Assembly-Directed Antivirals. J. Virol. (2018) doi:10.1128/jvi.01082-18.

57. Zlotnick, A. et al. Core protein: A pleiotropic keystone in the HBV lifecycle. Antiviral Research (2015) doi:10.1016/j.antiviral.2015.06.020.

58. Sclavi, B., Woodson, S., Sullivan, M., Chance, M. R. & Brenowitz, M. Time-resolved synchrotron x-ray ‘footprinting’, a new approach to the study of nucleic acid structure and function: Application to Protein-DNA interactions and RNA folding. J. Mol. Biol. (1997) doi:10.1006/jmbi.1996.0775.

59. Chandler-Bostock, R. et al. RNA X-ray footprinting reveals the consequences of an in vivo acquired determinant of viral infectivity. bioRxiv 2021.08.10.455819 (2021) doi:10.1101/2021.08.10.455819.

60. Jain, R. et al. New high-throughput endstation to accelerate the experimental optimization pipeline for synchrotron X-ray footprinting. J. Synchrotron Radiat. 28, 28 (2021).

61. Chandler-Bostock, R.; Bingham R.; Clark, S.; Scott, A. P.; Wroblewski, E.; Barker, A.; White, S. J.; Dykeman, E. J.; Mata, C. P.; Bohon, J.; Farquhar, E.; Twarock, R.; Stockley, P. G. RNA X-ray footprinting reveals an in vivo determinant of viral infectivity. Submitt. to Nat. Methods.

62. Oropeza, C. E. & McLachlan, A. Complementarity between epsilon and phi sequences in pregenomic RNA influences hepatitis B virus replication efficiency. Virology 359, 371–381 (2007).

63. Tang, H. & McLachlan, A. A pregenomic RNA sequence adjacent to DR1 and complementary to epsilon influences hepatitis B virus replication efficiency. Virology 303, 199–210 (2002).

64. Patel, N. et al. Dysregulation of Hepatitis B Virus Nucleocapsid Assembly with RNA-directed Small Ligands. bioRxiv 2021.08.10.455820 (2021) doi:10.1101/2021.08.10.455820.

65. Tillmann, H. L. et al. Safety and efficacy of lamivudine in patients with severe acute or fulminant hepatitis B, a multicenter experience. J. Viral Hepat. (2006) doi:10.1111/j.1365-2893.2005.00695.x.

66. Ying, C., De Clercq, E. & Neyts, J. Lamivudine, adefovir and tenofovir exhibit long-lasting anti-hepatitis B virus activity in cell culture. J. Viral Hepat. (2000) doi:10.1046/j.1365-2893.2000.00192.x.

67. Greenberg HB, Pollard RB, Lutwick LI, Gregory PB, Robinson WS, M. T. Effect of human leukocyte interferon on hepatitis B virus infection in patients with chronic active hepatitis. N Engl J Med. 295, 517–22 (1976).

68. Wu, S. et al. Discovery and Mechanistic Study of Benzamide Derivatives That Modulate Hepatitis B Virus Capsid Assembly. J. Virol. (2017) doi:10.1128/jvi.00519-17.

69. Venkatakrishnan, B. et al. Hepatitis B Virus Capsids Have Diverse Structural Responses to Small-Molecule Ligands Bound to the Heteroaryldihydropyrimidine Pocket. J. Virol. (2016) doi:10.1128/jvi.03058-15.

70. Zhang, X. et al. Discovery of Novel Hepatitis B Virus Nucleocapsid Assembly Inhibitors. ACS Infectious Diseases (2018) doi:10.1021/acsinfecdis.8b00269.

71. Schlicksup, C. J. et al. Hepatitis B virus core protein allosteric modulators can distort and disrupt intact capsids. Elife (2018) doi:10.7554/eLife.31473.

72. Yang, L. et al. Effect of a hepatitis B virus inhibitor, NZ-4, on capsid formation. Antiviral Res. (2016) doi:10.1016/j.antiviral.2015.11.004.

73. Stray, S. J. & Zlotnick, A. BAY 41-4109 has multiple effects on Hepatitis B virus capsid assembly. J. Mol. Recognit. (2006) doi:10.1002/jmr.801.

74. Stray, S. J. et al. A heteroaryldihydropyrimidine activates and can misdirect hepatitis B virus capsid assembly. Proc. Natl. Acad. Sci. U. S. A. (2005) doi:10.1073/pnas.0409732102.

75. Fatehi, F. et al. An intracellular model of hepatitis b viral infection: An in silico platform for comparing therapeutic strategies. Viruses 13, (2021).

76. Fatehi, F., Bingham, R. J., Dykeman, E. C., Stockley, P. G. & Twarock, R. An age-structured model of hepatitis B viral infection highlights the potential of different therapeutic strategies. (2021).

77. Strick-Marchand, H. et al. A novel mouse model for stable engraftment of a human immune system and human hepatocytes. PLoS One (2015) doi:10.1371/journal.pone.0119820.

78. Porterfield, J. Z. & Zlotnick, A. A simple and general method for determining the protein and nucleic acid content of viruses by UV absorbance. Virology 407, 281–288 (2010).

79. Scheres, S. H. W. RELION: Implementation of a Bayesian approach to cryo-EM structure determination. J. Struct. Biol. (2012) doi:10.1016/j.jsb.2012.09.006.

80. Zivanov, J. et al. New tools for automated high-resolution cryo-EM structure determination in RELION-3. Elife (2018) doi:10.7554/eLife.42166.

81. Zheng, S. Q. et al. MotionCor2: Anisotropic correction of beam-induced motion for improved cryo-electron microscopy. Nature Methods (2017) doi:10.1038/nmeth.4193.

82. Zhang, K. Gctf: Real-time CTF determination and correction. J. Struct. Biol. (2016) doi:10.1016/j.jsb.2015.11.003.

83. Pettersen, E. F. et al. UCSF Chimera - A visualization system for exploratory research and analysis. J. Comput. Chem. (2004) doi:10.1002/jcc.20084.

84. Adams, P. D. et al. PHENIX: A comprehensive Python-based system for macromolecular structure solution. Acta Crystallogr. Sect. D Biol. Crystallogr. (2010) doi:10.1107/S0907444909052925.

85. Emsley, P. & Cowtan, K. Coot: Model-building tools for molecular graphics. Acta Crystallogr. Sect. D Biol. Crystallogr. (2004) doi:10.1107/S0907444904019158.

86. Chen, V. B. et al. MolProbity: All-atom structure validation for macromolecular crystallography. Acta Crystallogr. Sect. D Biol. Crystallogr. (2010) doi:10.1107/S0907444909042073.

## References

1. Pollack, J. R. & Ganem, D. An RNA stem-loop structure directs hepatitis B virus genomic RNA encapsidation. J. Virol. (1993) doi:10.1128/jvi.67.6.3254-3263.1993.

2. Junker-Niepmann, M., Bartenschlager, R. & Schaller, H. A short cis-acting sequence is required for hepatitis B virus pregenome encapsidation and sufficient for packaging of foreign RNA. EMBO J. (1990) doi:10.1002/j.1460-2075.1990.tb07540.x.

3. Ding, Y., Chan, C. Y. & Lawrence, C. E. Sfold web server for statistical folding and rational design of nucleic acids. Nucleic Acids Res. (2004) doi:10.1093/nar/gkh449.

4. Patel, N. et al. HBV RNA pre-genome encodes specific motifs that mediate interactions with the viral core protein that promote nucleocapsid assembly. Nat. Microbiol. 2, 17098 (2017).

